# Perturb-map enables CRISPR genomics with spatial resolution and identifies regulators of tumor immune composition

**DOI:** 10.1101/2021.07.13.451021

**Authors:** Maxime Dhainaut, Samuel A Rose, Guray Akturk, Aleksandra Wroblewska, Eun Sook Park, Sebastian R Nielsen, Mark Buckup, Vladimir Roudko, Luisanna Pia, Jessica Le Berichel, Anela Bektesevic, Brian H Lee, Alessia Baccarini, Nina Bhardwaj, Adeeb H Rahman, Sacha Gnjatic, Dana Pe’er, Miriam Merad, Brian D Brown

**Affiliations:** Precision Immunology Institute, Icahn School of Medicine at Mount Sinai, New York, NY; Icahn Genomics Institute, Icahn School of Medicine at Mount Sinai, New York, NY; Tisch Cancer Institute, Icahn School of Medicine at Mount Sinai, New York, NY, USA; Department of Genetics and Genomic Sciences, Icahn School of Medicine at Mount Sinai, New York, NY; Computational and Systems Biology Program, Memorial Sloan Kettering Cancer Center, New York, NY

## Abstract

The cellular architecture of a tumor, particularly immune composition, has a major impact on cancer outcome, and thus there is an interest in identifying genes that control the tumor microenvironment (TME). While CRISPR screens are helping uncover genes regulating many cell-intrinsic processes, existing approaches are suboptimal for identifying gene functions operating extracellularly or within a tissue context. To address this, we developed an approach for spatial functional genomics called Perturb-map, which utilizes protein barcodes (Pro-Code) to enable spatial detection of barcoded cells within tissue. We show >120 Pro-Codes can be imaged within a tumor, facilitating spatial mapping of 100s of cancer clones. We applied Perturb-map to knockout dozens of genes in parallel in a mouse model of lung cancer and simultaneously assessed how each knockout influenced tumor growth, histopathology, and immune composition. Additionally, we paired Perturb-map and spatial transcriptomics for unbiased molecular analysis of Pro-Code/CRISPR lesions. Our studies found in Tgfbr2 knockout lesions, the TME was converted to a mucinous state and T-cells excluded, which was concomitant with increased TGFβ expression and pathway activation, suggesting Tgfbr2 loss on lung cancer cells enhanced suppressive effects of TGFβ on the TME. These studies establish Perturb-map for functional genomics within a tissue at single cell-resolution with spatial architecture preserved.

## INTRODUCTION

The cellular composition, spatial architecture, and tissue localization of a tumor have a major impact on a cancer’s progression and response to therapy. These factors are particularly relevant to tumor immunity as immune cell composition and localization within the tumor microenvironment (TME) is one of the major determinants of response to immunotherapy and patient outcome (Binnewies et al., 2018; Chen and Mellman, 2017; Cristescu et al., 2018; Riaz et al., 2017; Taube et al., 2018). The general view is that immunosuppressive macrophages and regulatory T cells are recruited into tumors and effector CD8 T cells excluded, and this prevents cancer cell killing. Fibroblasts in the tumor stroma, depending on their activation state, can also prevent T cell infiltration and killing by remodeling the TME and helping to create an immunosuppressive state (Turley et al., 2015). Immune exclusion can be highly local, often occurring just in small regions of the tumor (Keren et al., 2018), which represent potential pockets of immune resistance. The mediators of this heterogeneity in tumor composition are not known. Spatially resolved single cell genome sequencing has revealed that many genetically distinct sub-clones exist in proximity in tumors (Minussi et al., 2021), but how different mutations influence the TME (Galluzzi et al., 2018; Mitra et al., 2020), or how neighboring clones influence one another is not well-defined. Indeed, while some genes involved in orchestrating the TME have been identified, the potential role of many genes in influencing the architecture and immune composition of different tumors are not established.

Identifying the genes controlling the cellular arrangement of tumors is a challenge. TME composition is complex; comprised of many different immune and non-immune cell types whose ratio, location, and movement are all interdependent factors (Binnewies et al., 2018). Many gene’s functions are dependent on the context of a tissue and spatial proximity to specific cell types and structures (Haigis et al., 2019). For example, T cell migration from tissues to lymph nodes is dependent on the chemokines CCL19 and CCL21 (Hauser and Legler, 2016). The functions of these genes, and their receptor CCR7, only fully emerges in the context of a discrete tissue and organ arrangement that includes lymphatics and vasculature. This is not unique to immune cells or chemokines and applies to many classes of genes. Thus, determining which of the many 100s of genes expressed in a tumor influence TME composition and arrangement ultimately requires in vivo studies. However, knockout (KO) or overexpression (OE) of 100s of genes in separate animal models is not practically feasible.

To scale up studies of gene functions, pooled CRISPR screens are increasingly being used (Doench, 2018). Cells can be transduced with 100s of CRISPR vectors, and the frequency of cells carrying each vector determined using the CRISPR as a ‘barcode’. Gene function is inferred by applying a selective pressure, such as time or a drug, and measuring changes in CRISPR frequency (Shalem et al., 2015). Single cell sequencing approaches, such as Perturb-seq and ECCITE-seq, and high-dimensional cytometry, have further advanced CRISPR screens by enabling the molecular and phenotypic changes caused by gene perturbation to be measured more comprehensively (Adamson et al., 2016; Dixit et al., 2016; Jaitin et al., 2016; Mimitou et al., 2019; Wroblewska et al., 2018). CRISPR screens have been utilized to identify genes involved in cancer sensitivity to immune editing, and helped establish a number of important regulators of this process, such as Ptpn2 and the IFNγ pathway (Lawson et al., 2020; Manguso et al., 2017; Patel et al., 2017).

While pooled approaches enable scaled throughput, existing technologies have limitations for in vivo studies. One of the most significant is that tissue is dissociated for analysis. This largely restricts the biological functions that can be probed with pooled screens to cell intrinsic processes, as the extracellular effects of a gene perturbation cannot be assessed once tissues are homogenized. This excludes using CRISPR genomics to identify genes controlling phenotypes that require spatial resolution to assess, such as immune cell localization or vascular density within a tumor. In addition, gene functions that are mediated extrinsically can potentially be compensated by adjacent cells that do not carry the same KO. Thus, with current pooled CRISPR approaches it is not readily feasible to determine the functions of a secreted factor, such as a chemokine or interleukin, or a regulator of one of these factors, as the local effects from KO in a small fraction of cells cannot be measured or could be counteracted by cells with a normal copy of the same gene. Recently, a novel approach was described for detecting barcoded CRISPR vectors by imaging, which was shown to enable assaying of pooled screens without cell disruption(Feldman et al., 2019). This is a very powerful high-resolution technology but has not yet been used for tissue level analysis.

Here, we describe an approach for in vivo spatial functional genomics, called Perturb-map. Perturb-map is based on a protein barcode (Pro-Code) system that utilizes triplet combinations of a small number of linear epitopes to create a higher order set of unique barcodes that can mark cells expressing different CRISPR gRNAs (Wroblewska et al., 2018). We show that >120 different Pro-Code expressing cancer cell populations can be detected within a tumor at single cell resolution and tissue scale. We applied Perturb-map to KO 35 genes in parallel in a mouse model of lung cancer and simultaneously assessed how each KO influenced key parameters of tumor biology, including growth, histopathology, and immune composition. We also paired Perturb-map with spatial transcriptomics to provide a broad analysis of the molecular state of different gene-targeted tumor lesions. Amongst our findings, we observed that Socs1 loss in lung cancer cells provided a growth advantage but was accompanied by T cell recruitment into the tumors, while knockout of Tgfbr2 also resulted in a growth advantage but lead to conversion of the TME to a mucinous state and resulted in T cell exclusion from the TME. A striking finding, which was revealed by our ability to assess different gene perturbations in parallel in situ, was how spatially segregated the effects of Socs1 and Tgfbr2 KO were, as T cell infiltration and exclusion were tightly confined to the Socs1 and Tgfbr2 lesions, respectively, even when the two were in adjacent proximity.

These studies establish the use of Perturb-map for broad phenotypic analysis of dozens of genes in parallel within a tissue or tumor at cellular resolution with spatial architecture preserved.

## RESULTS

### In situ detection of 120 Pro-Code populations in lung and breast tumors by multiplex imaging

In previous studies, we described a novel protein-based vector/cell barcoding system, the Pro-Codes, which is comprised of triplet combinations of linear epitopes (e.g. FLAG, HA, AU1, etc.) fused to a scaffold protein, dNGFR (Wroblewska et al., 2018). As the Pro-Codes are detected by antibody staining, we hypothesized that they could be resolved by imaging. A number of techniques have been developed for multiplex staining of tissues for histological analysis (Gut et al., 2018; Lin et al., 2018; Remark et al., 2016; Tsujikawa et al., 2017). The general principle of these approaches involves staining sections with 1 - 4 antibodies, imaging, stripping, then re-staining. As many as 60 markers can be detected in a single section (Lin et al., 2018). Our group previously developed a variant of this approach termed multiplex immunohistochemistry consecutive staining on a single slide (MICSSS) (Remark et al., 2016).

To test detection of the Pro-Codes by imaging, we first generated cancer lines expressing the Pro-Codes. We transduced mouse Kras*^G12D^* p53^−/−^ (KP) lung cancer cells (DuPage et al., 2009) and 4T1 breast cancer cells (4T1) with a pool of lentiviral vectors (LV) encoding 84 or 120 different Pro-Codes. We injected mice i.v. with KP cells or into the mammary fad pad for 4T1 cells. After 2 weeks (4T1) or 4 weeks (KP), the tumor bearing tissue was collected, fixed, embedded in paraffin, sectioned, and stained for each of the Pro-Code epitopes using MICSSS (**Figure S1A**). The images were registered, deconvoluted (to separate the hematoxylin and epitope signal) and overlaid to visualize epitope tag colocalization. Alternatively, tissue sections were stained with metal-conjugated antibodies and imaged on a multiplex ion beam imager (MIBI) (**Figure S1A and B**). Each epitope could be efficiently detected with both techniques at a sub-cellular resolution, as we could detect the epitopes at the cell membrane, as expected from the membrane-localizing dNGFR scaffold. We were therefore able to spatially resolve up to 120 Pro-Codes in breast (**Figure 1A and B**) and lung (**Figure 1C**) tumor models. This is >10-fold the number of spatially resolvable reporters used for comparable system, such as confetti mice (Schepers et al., 2012), and, to our knowledge, represents the largest cell barcoding system detectable by antibody-based histological imaging.

**Figure 1.**
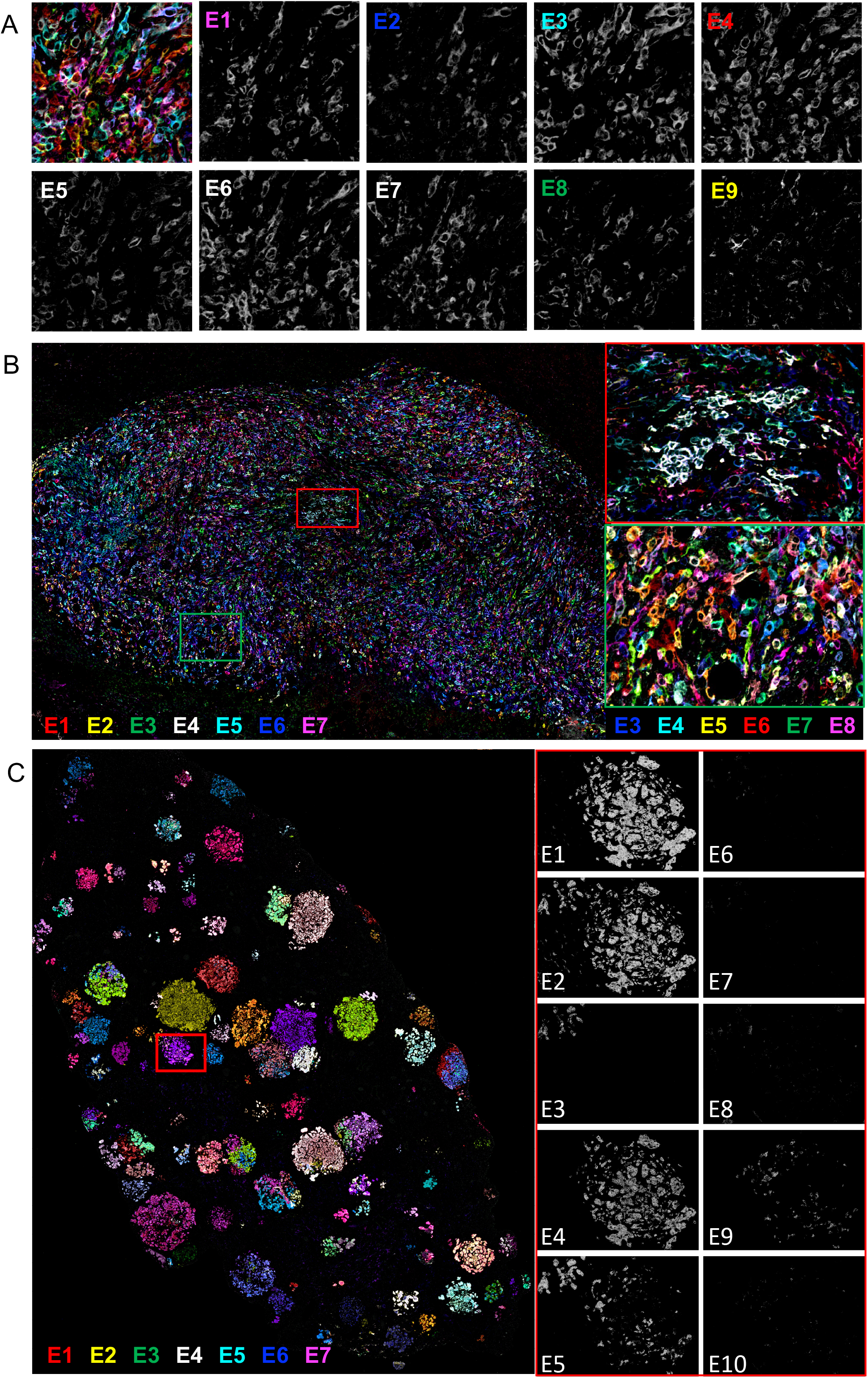
Multiplex imaging maps Pro-Code labeled breast and lung cancer populations in breast and lung tissue sections. (**A**) Pro-Code epitope tag detection in 4T1 breast tumors. 4T1 mammary carcinoma cells were transduced with a dNGFR-Pro-Code (memPC) library (84 or 120 memPC with 9 or 10 epitope tags, respectively) and injected into the mammary fat pad of female BALB/c mice. The tumors were harvested 14 days later, formalin-fixed paraffin-embedded (FFPE), and stained using MICSSS. Representative images showing detection of 9 epitope tags. Panel 1 (upper left corner) shows an overlay of tags E1, E2, E3, E4, E8, and E9. (**B**) Imaging of Pro-Code expressing 4T1 breast tumors. (Left) Shown is a representative whole tumor image of a memPC-expressing 4T1 breast tumor generated as described in (A). In the image, 7 epitope tags are represented simultaneously, color-coded as indicated. (Right) Zoomed-in images of the highlighted regions (red and green rectangle) of the same tumor, displaying respectively a clonal and mixed distribution of 4T1 cells. Images are representative of more than 10 tumors (n=10 mice), across 2 different experiments with 84 or 120 different memPC. (**C**) Imaging of Pro-Code expressing KP lung adenocarcinomas in mouse lungs. KP lung cancer cells were transduced with a memPC library (120 memPC, 10 epitope tags) and injected i.v. in C57BL/6 mice. The lungs were harvested 4 weeks later, fixed, paraffin-embedded and stained using MCISSS. (Left) Representative overlaid image of a whole lobe of lung with memPC expressing KP tumor lesions. 7 epitope tags are represented simultaneously, color-coded as indicated. (Right) Zoomed-in images corresponding to the highlighted region (red rectangle), displaying individual epitope tag stainings. Images representative of 10 mice (5 lobes per mouse), across 2 different experiments with different memPC libraries.

Though we could readily detect each of the Pro-Code epitopes, membrane stains are still not optimal for cell segmentation, which makes it more difficult for downstream analysis. To address this, we created a new set of Pro-Codes which utilized a nuclear localizing mCherry fluorescent protein (mCherry-NLS) as a scaffold. Sequences encoding 165 triplet combinations of 11 epitope tags were cloned in frame into the N-terminal domain of mCherry-NLS within the LV backbone to create a nuclear Pro-Code (nPC) library. We transduced 293T cells with the library of 165 nPC, after 5 days we performed intracellular staining for each of the 11 epitopes, and analyzed by CyTOF. Similar to the membrane-bound Pro-Codes (memPC), we could detect each of the nPC by CyTOF with single cell resolution (**Figure S1C and S1D**). Next, we transduced 4T1 and KP cells with a library of 120 nPC and repeated the experiments described above, using MICSSS for Pro-Code detection (**Figure S1A**). We were able to detect each of the 120 unique nPC in both the lung and breast tumor models, and found that they localized to the nucleus (**Figure 2A**).

**Figure 2.**
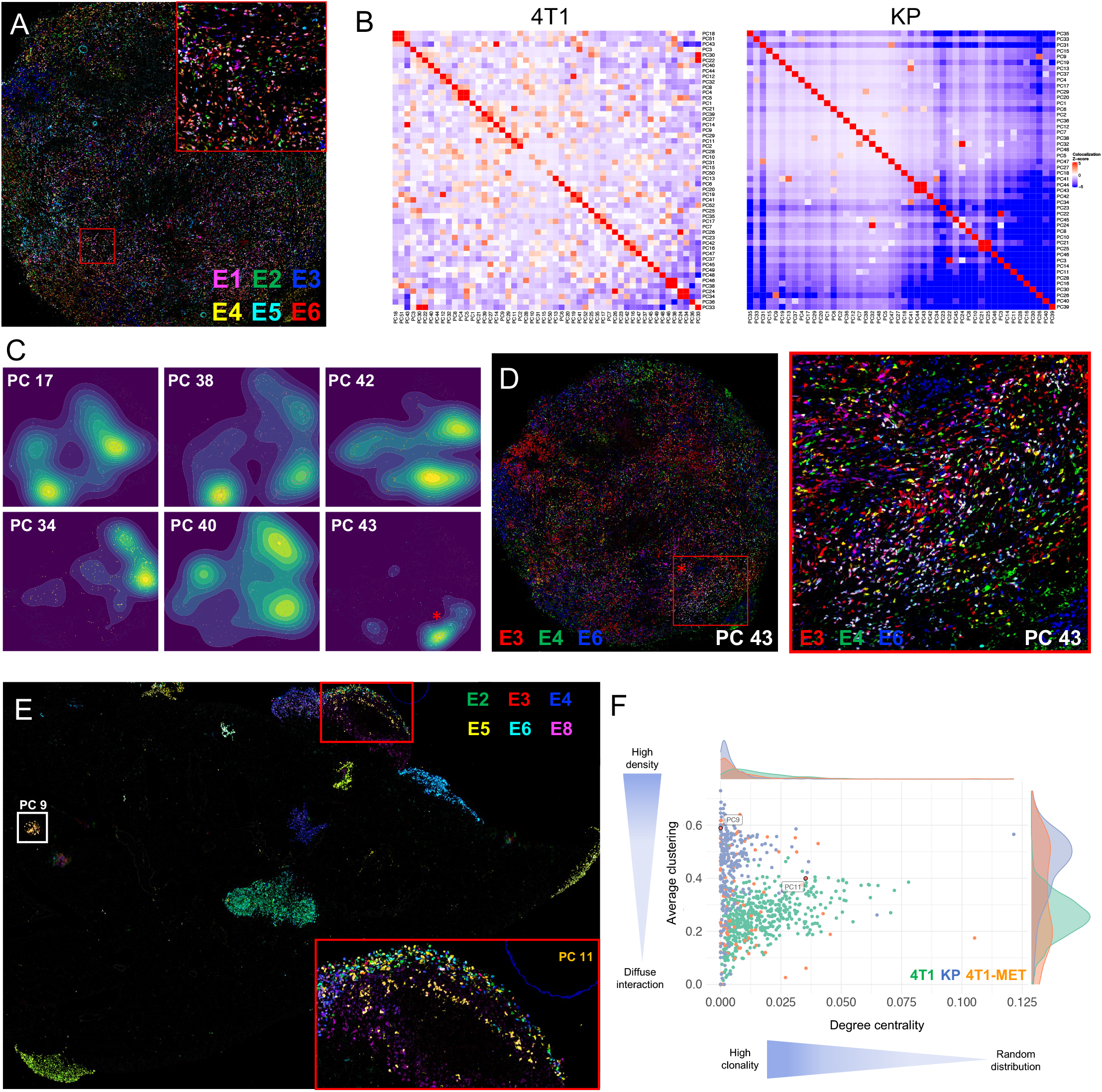
Analysis of tumor development by Pro-Code mapping reveals tumor clonal heterogeneity. (**A**) Imaging of nuclear Pro-Code expressing breast tumors. 4T1 cells were transduced with a nuclear Pro-Code (nPC) library (120 nPC, 10 epitope tags) and injected into the mammary fat pad of female BALB/c mice. The primary tumors were harvested 14 days later, tissues were formalin fixed paraffin-embedded (FFPE) and stained using MICSSS. Shown is a representative image of a tumor with six epitope tags represented, color-coded as indicated. The highlighted region (red square) is enhanced on the top right and shows nuclear localization of the nPC. Image representative of 16 tumors (n=16 mice), across 2 different experiments with two different nPC libraries. (**B**) Colocalization analysis quantifying the relative frequency of interactions between nPC populations in a breast tumor (left) or lung tumor lesions (right). Pro-Code expressing 4T1 breast and KP lung tumors were generated and imaged as described above and in Figure 1. Cell-to-cell interactions were quantified on a neighbors graph constructed using the 10 nearest neighbors at a maximum radius of 40 mm from each cell. Each square represents interactions between two PC populations and is shaded based on the significance (Z-Score) if the observed interaction frequency relative to a permuted background distribution generated by swapping Pro-Code labels (1,000 permutations). (**C**) Density maps of 6 randomly selected nPC populations from the 4T1 breast tumor displayed in Figure 2D. Kernel density estimates were generated using nPC cell coordinates with a bivariate normal kernel and bandwidth determined by normal distribution approximation. The star corresponds to the example highlighted in Figure 2D. (**D**) Overlayed image of the 3 epitope tags corresponding to Pro-Code 43 (PC 43), color-coded as indicated. Cells expressing the 3 epitope tags (i.e. PC 43) appear white. On the right is a Zoomed-in image of the highlighted region (red square). (**E**) Imaging of nPC labelled 4T1 breast metastases in the lung. nPC-expressing 4T1 cells (120 nPC library) were injected into the mammary fat pad of BALB/c mice. Lungs were harvested after 28 days and FFPE sections stained for Pro-Code epitopes by MICSSS. Six epitope tags are represented, color-coded as indicated. PC 9 (white square) and PC 11 (gold cells in the left square) are highlighted. (**F**) Comparative analysis of tumor development of 4T1 primary breast tumors in the mammary fatpad (green), KP tumor lesions in the lung (blue), and 4T1 metastases in the lung (orange). Plotted are the average local clustering coefficient and group degree centrality (fraction of alternate Pro-Code neighbors) of each Pro-Code population within a neighbors graph constructed on Pro-Code positive cells (k = 10, maximum cell radius 40 mm). Each point represents one Pro-Code on an individual tissue section. Only Pro-Codes present in >20 cells on a tissue section are displayed. Data based on mice analyzed from above (A), (E), and Figure 1.

Since the memPC and nPC have a distinct subcellular localization, we hypothesized that we could use them both in combination in the same cells. We transduced 4T1 cells with a library of 56 memPC (8 tags), sorted the dNGFR positive cells, and further transduced them with a library of 56 nPC that had the same 8 tags. The cells were injected into mice, as above (**Figure S1E**). In the resulting tumors, we were able to detect both mCherry and NGFR on the same cells. More importantly, we were able to identify cells expressing a different Pro-Code in the nucleus and at the cell membrane, exponentially increasing the amount of spatially resolvable reporters for cell tracking purposes to up to 3,136 combinations with only 8 epitope tags (**Figure S1F and S1G**).

### Pro-Code imaging reveals KP lung and 4T1 breast tumor clonality

An immediately apparent difference revealed by imaging the Pro-Codes in the two tumor types was the highly heterogeneous distribution of Pro-Codes in 4T1 tumors compared to KP lung tumors, in which almost all of the tumor lesions were clonal, as evident by being positive for only 3 epitopes corresponding to a specific Pro-Code (**Figure 1B,C** and **S1B**). This implied that each KP tumor lesion is initiated by a single KP cancer cell. Moreover, as the tumor burden increased and tumor lesions developed in close proximity, there appeared to be minimal mixing between adjacent lesions (**Figure 1C**).

To further analyze the clonal dynamics of 4T1 and KP tumors, we performed nuclear segmentation of the overlaid image based on the hematoxylin signal. The resulting table, in which each cell is represented by a set of coordinates in 2D space along with the expression intensity of each epitope tag, was used to debarcode the cells (i.e. identify expressed Pro-Code) using an adapted form of a debarcoding algorithm used for the analysis of cell suspensions (Zunder et al., 2015). This allowed us to reconstruct the image and pseudo-color each cell based on the Pro-Code they express, facilitating all downstream analyses (**Figure S2A and B**). Next, we assessed the co-localization between Pro-Code populations within 4T1 primary tumors and KP tumor lesions (**Figure 2B**). For each Pro-Code population, a Z-score was calculated based on the number of observed interactions on a nearest neighbor graph between itself and all Pro-Code populations on the tissue section relative to a permuted null distribution generated by randomly swapping cells. This confirmed that KP tumors were highly clonal, as each Pro-Code population mainly exhibited homotypic interactions. Surprisingly, although the 4T1 cells appeared to be randomly distributed, co-localization analysis also showed a certain degree of clonality, although with an increased level of heterotypic interactions compared to the KP tumors. Mapping of each Pro-Code within the primary tumor confirmed that cells expressing a given Pro-Code tended to cluster together (**Figure S2C**).

To better understand how the regional clusters were forming, we visualized the relative regional abundance of Pro-Code-positive cells with a bivariate normal kernel density estimation (**Figure 2C**). This revealed that cells within the primary 4T1 lesion spread from one or several foci, with the cell density decreasing as the distance from the focal point increases. Although this phenomenon was not apparent on complex overlaid images (**Figure 1B and 2A**), it became apparent when we only represented 3 epitope tags corresponding to a given Pro-Code. For example, our analysis revealed that cells expressing PC43 mainly clustered in one location (**Figure 2C and 2D**). This indicates a diffuse clonality characteristic of 4T1 in the primary tumor and supports a model in which 4T1 cancer cells migrate locally from a focal point of origin.

Consistent with their high motility, 4T1 breast tumors are prone to metastasis (Tao et al., 2008). We sought to use the Pro-Codes to assess the potential clonal heterogeneity of 4T1 tumor derived metastasis. We injected nPC-expressing 4T1 cells into the mammary fat pad to generate breast tumors. After 4 weeks, to allow for lung metastasis, we collected the lungs, and stained sections for the Pro-Codes by MICSSS (**Figure 2E**). Many of the metastasis were homogeneous for a single Pro-Code (e.g. PC9), indicating they had originated from a single cell from the primary tumor. However, there were also metastatic lesions in the lung that contained a mix of Pro-Code expressing cells, suggesting either multi-cell seeding or that initial seeding by a clone was followed by additional metastatic cells. We compared the clonal heterogeneity of all three models (KP lung, 4T1 breast, 4T1 breast metastasis in the lung) by assessment of average clustering coefficient and the group degree of centrality (fraction of non-group neighbors) for each Pro-Code population (**Figure 2F**). This confirmed the distinct spatial patterning of each tumor context, with KP lung tumors having high clonality and low mixing, 4T1 primary tumors having diffuse clonality, and 4T1 metastases in the lung developing clonally, but also with a bimodal pattern in the density of interactions, reflecting the presence of mixed metastases. They also demonstrate how multiplex imaging of the Pro-Codes (Pro-Code-map) can be used to reveal tumor clonality and heterogeneity within a tissue.

### Perturb-map identifies regulators of sensitivity to immunoediting and tumor architecture

Cancer immunotherapy, particularly checkpoint blockade, has become a highly effective treatment for some cancers. However, many patients only have a partial or even no response to therapy, and thus there is a significant interest in identifying factors that control tumor immunity(Sharma et al., 2021). CRISPR screens have helped identify some of the genes involved, but as noted, current approaches are not suited to study many key aspects of tumor immunology, particularly immune cell recruitment and exclusion. Having established we could detect the Pro-Codes by imaging, we set out to determine if we could use the Pro-Codes to create a platform for resolving CRISPR screens in situ, and specifically to enable us to determine how different immunomodulatory proteins might influence the immune cell composition of a tumor.

We built a library of nPC/CRISPR LV vectors targeting 35 genes coding for regulators of cytokine signaling pathways (including receptors for IFNα, IFNγ, TGFβ, and TNFα) and ligands and secreted factors involved in immune cell interactions (e.g. B2m, Cd47, Cd247, and Cxcl17) (**Figure 3A**). Though these genes are known to have immune functions, their roles in lung TME biology are not fully established. As a control, we targeted the F8 gene, which is not expressed by KP cells. For each of the 35 genes, KP cells encoding Cas9 were transduced with 3 different LV, with each vector expressing the same nPC but a different gRNA targeting the same gene. The 35 populations of nPC/CRISPR KP were then pooled in equal proportions by cell sorting. The frequency of each nPC/CRISPR population was determined by CyTOF and confirmed to be in similar distribution. We injected the cells i.v. into Cas9-expressing mice (n=11) to seed tumors and after 4 weeks we collected the lungs for tumor analysis (**Figure 3B**). Tissues were sectioned and stained for the Pro-Code by MICSSS. MICSSS was used because it preserves tissue architecture and is compatible with whole slide scanning, so that all lung lobes from a mouse could be imaged together at high resolution.

**Figure 3.**
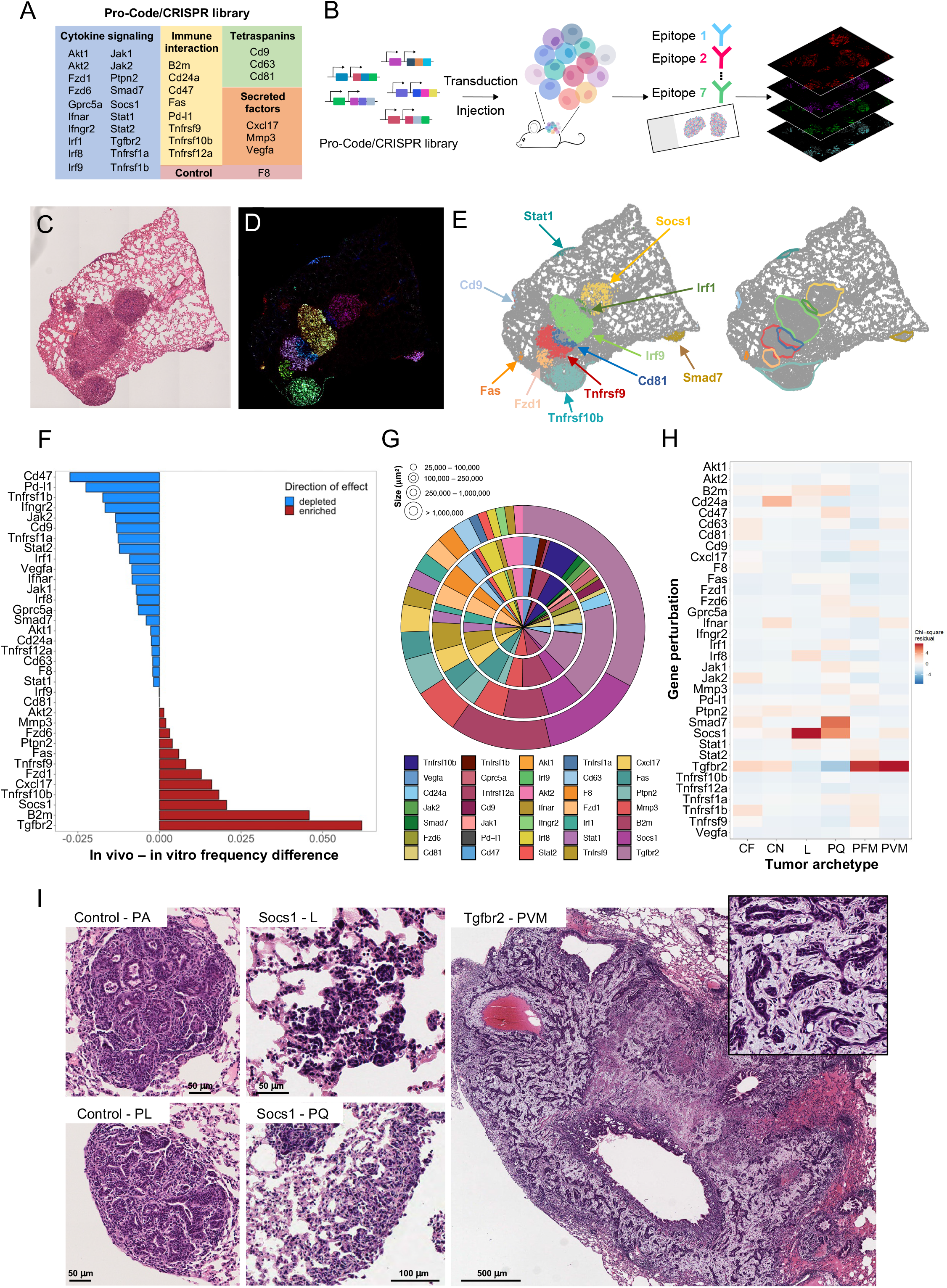
Perturb-map identifies genes regulating tumor development and architecture in vivo. (**A**) Table of the genes targeted in the Pro-Code/CRISPR library. (**B**) Schematic of the Perturb-map experimental pipeline. For the studies in Figures 3 and 4, KP-Cas9 cells were transduced with the nPC/CRISPR library described in Figure 3A and injected into 11 mice from 2 independent experiments (n=5 and n=6). Lungs were collected after 4 weeks, and formalin-fixed paraffin-embedded (FFPE) tissue sections were stained using by MICSSS for 7 different epitope tags to detect 35 Pro-Code populations and for specific cell markers (see Figure 4). Stained sections were imaged by whole slide scanning. (**C-E**) Overview of the Perturb-map analysis pipeline. (**C**) Representative example of an H&E stained tissue section. Shown is a section of 1 lung lobe from a mouse injected i.v. with KP-Cas9 cells transduced with the nPC/CRISPR library from (A). (**D**) A serial section from the same tissue as in (C) was stained for all 7 epitope tags by MICSSS. The image represents an overlay of pseudo-colored epitope tag stainings. Different colored areas correspond to different Pro-Code populations. (**E**) Debarcoded and reconstituted digital image of the tissue section in D. (Left panel) Each dot represents a cell, colored based on Pro-Code expression (Pro-Code negative cells in grey). Gene perturbations can be annotated directly on the image based on Pro-Code expression. (Right panel) Tumor boundaries were defined following DBSCAN clustering of Pro-Code positive cells. (**F**) Relative frequency of Pro-Code/CRISPR KP tumors. The frequency of each Pro-Code/CRISPR KP-Cas9 population was determined pre-injection (by CyTOF) and compared to the relative abundance of corresponding Pro-Code/CRISPR lesions in vivo. Deviations from 0 indicate positive or negative selection for each Pro-Code/CRISPR population *in vivo*. (**G**) Quantification of tumor lesions size across gene perturbations. Shown are the percentages of tumors associated with each gene perturbation within discrete tumor size categories. Each ring corresponds to a tumor size range, as indicated. (**H**) Histopathology analysis of Pro-Code/CRISPR tumor lesions. A total of ∼1,750 tumors were scored by a pathologist on H&E sections (with no Pro-Code staining or identification markers) and tumor archetype identified. The heatmap shows the standardized residuals of a Chi-squared test between gene perturbation and tumor archetype to identify gene-arcehtype associations (Chi-squared p value 4.43×10^−14^). (**I**) Representative example of tumor archetypes associated with Socs1 and Tgfbr2 gene perturbation. PA: parenchymal, PL: pleural, L: lepidic, PQ: pleural plaque, PVM: perivascular mucinous tumor.

For each sample, we stained serial sections with hematoxylin and eosin or with antibodies specific for the Pro-Code epitope tags (**Figure 3C and 3D**). The images were then segmented and debarcoded to identify the Pro-Codes expressed by each tumor lesion (**Figure 3E**, left panel). We used the density-based spatial clustering of applications with noise (DBSCAN) algorithm to sub-cluster Pro-Code populations into discrete lesions and infer tumor boundaries using alpha shapes (**Figure 3E**, right panel). We then used the Pro-Code identification to determine which CRISPR gRNAs were expressed in each tumor lesion. From 11 mice injected with the nPC/CRISPR KP cells, we identified approximately 1,750 distinct Pro-Code-expressing tumor lesions.

To determine if any of the targeted genes impacted tumor development in vivo, we compared the proportion of tumors carrying a specific nPC/CRISPR to the relative frequency of the same nPC/CRISPR in the pre-injected cell mix (as determined by CyTOF) (**Figure 3F**). There was no change in the relative proportion of our control, F8, as well as several other targeted genes, but there were substantial changes in the frequency of a number of Pro-Code/CRISPR, inferring the associated genes influence tumor growth. Two of the most depleted gene targets were the immune checkpoints Cd274 (Pd-l1) and Cd47, indicating loss of these genes impaired KP growth and implying KP tumors subdue both innate and adaptive immune pathways for development. Conversely, there was a significant enrichment of B2m targeted lesions, indicating loss of MHC class I presentation facilitated tumor growth, which further signified a role for adaptive immune control.

In contrast to what has been reported in many in vitro CRISPR screens, including our own (Wroblewska et al., 2018), we observed a de-enrichment of PC/CRISPRs targeting positive regulators of IFNγ signaling (Ifngr2, Jak2, Irf1, Jak1), along with enrichment of CRISPR targeting Socs1, a negative regulator of IFNγ signaling. While this differs from in vitro and some in vivo findings, it is consistent with studies from the Minn lab which found that IFNγ signaling specifically on cancer cells can help them escape immune control (Benci et al., 2016, 2019). Of note, Tgfbr2 targeted lesions were the most enriched in vivo indicating loss of the TGFβ receptor on KP cells enhanced tumor growth. Tgfbr2 tumors were also amongst the largest tumors in area, along with Socs1 and B2m (**Figure 3G**), which were also amid the most enriched. Though frequency and size did not always correlate, as there were relatively few Ifngr2 targeted tumors, but the ones that did establish were larger in size (**Figure 3G**), which may relate to how these genes influence different aspects of tumor biology.

In addition to being able to measure the number and area of tumor lesions associated with a gene KO, Perturb-map also enabled us to assess how different genes affected tumor architecture. To do so, each lesion was scored by a pathologist (blinded to the perturbations) following a list of standard clinical criteria that included the differentiation degree of the cancer cells, the location of lesion and the composition of the stroma (**Figure S3A and S3B**). Distinct tumor archetypes were identified based on a combination of features, including central necrotic (CN) or central fibrotic (CF) tumors, poorly differentiated pleural plaque (PQ) tumors, fibro-mucinous stroma (PFM) tumors, lepidic tumors (L) that formed along the lining of the lung alveoli, and perivascular mucinous tumors (PVM) which presented a mucinous stroma and surrounded vasculature. We performed a Chi-squared test to identify significant associations between gene perturbations and histological characteristics of the lesions (**Figure 3H** and **Figure S3B and S3C**). Interestingly, gene perturbations of Socs1 and Tgfbr2 led to the development of markedly different KP lung tumor lesions. Loss of Socs1 correlated to the development of pleural plaques (PQ) and lepidic (L) tumors, two poorly differentiated tumor lesion archetypes. Loss of Tgfbr2 resulted in a remodeling of the stromal compartment and induced the development of highly mucinous (PFM and PVM) tumors (**Figure 3H and 3I** and **Figure S3B and S3C**). This was not related to tumor size or lesion number, as both Socs1 and Tgfbr2 targeted lesions were similar in this regard but had very distinct histological states.

These studies establish the ability of the Perturb-map approach to facilitate spatial mapping of CRISPR-expressing lesions on tissue sections and assess the influence of many genes on tumor biology in parallel. They also indicate that loss of Tgfbr2 and Socs1 alter KP lung tumor architectures, in profoundly different ways, but each confers a growth advantage.

### Perturb-map identifies genes modulating the immune composition of the TME

Since immune cell populations can be identified in situ by marker expression, we aimed to use Perturb-map to investigate how different gene perturbations impact immune cell recruitment and maintenance in and around tumor lesions. To do this, we stained lung tissue sections that were stained for the Pro-Codes with antibodies specific for T cell (CD4, CD8), B cell (B220), and myeloid cell (F4/80, CD11b, CD11c) markers, as well as for EpCAM, an adhesion molecule that marks epithelial cells and is highly expressed on KP cancer cells. Pro-Code debarcoding identified the border and gene perturbations associated with each tumor (**Figure 4A**). The coordinates of each immune cell type were then used to determine their relative position to each tumor lesion and to calculate the density of immune infiltrates in the different tumor lesions (**Figure 4B and 4C**). Additionally, we measured the mean EpCAM intensity within the tumor borders. Tumor lesions for each gene perturbations were then compared to control lesions (carrying CRISPR gRNAs targeting F8) using a Wilcoxon test. We excluded from the analysis any gene perturbation that was not found in at least 20 tumor lesions in vivo. The resulting Z-scores and p-values were represented in radial plots, which indicate the relative infiltration (outside facing bars) or exclusion (inside facing bars) of each immune cell type within the indicated Pro-Code/CRISPR lesions (**Figure 4D**).

**Figure 4.**
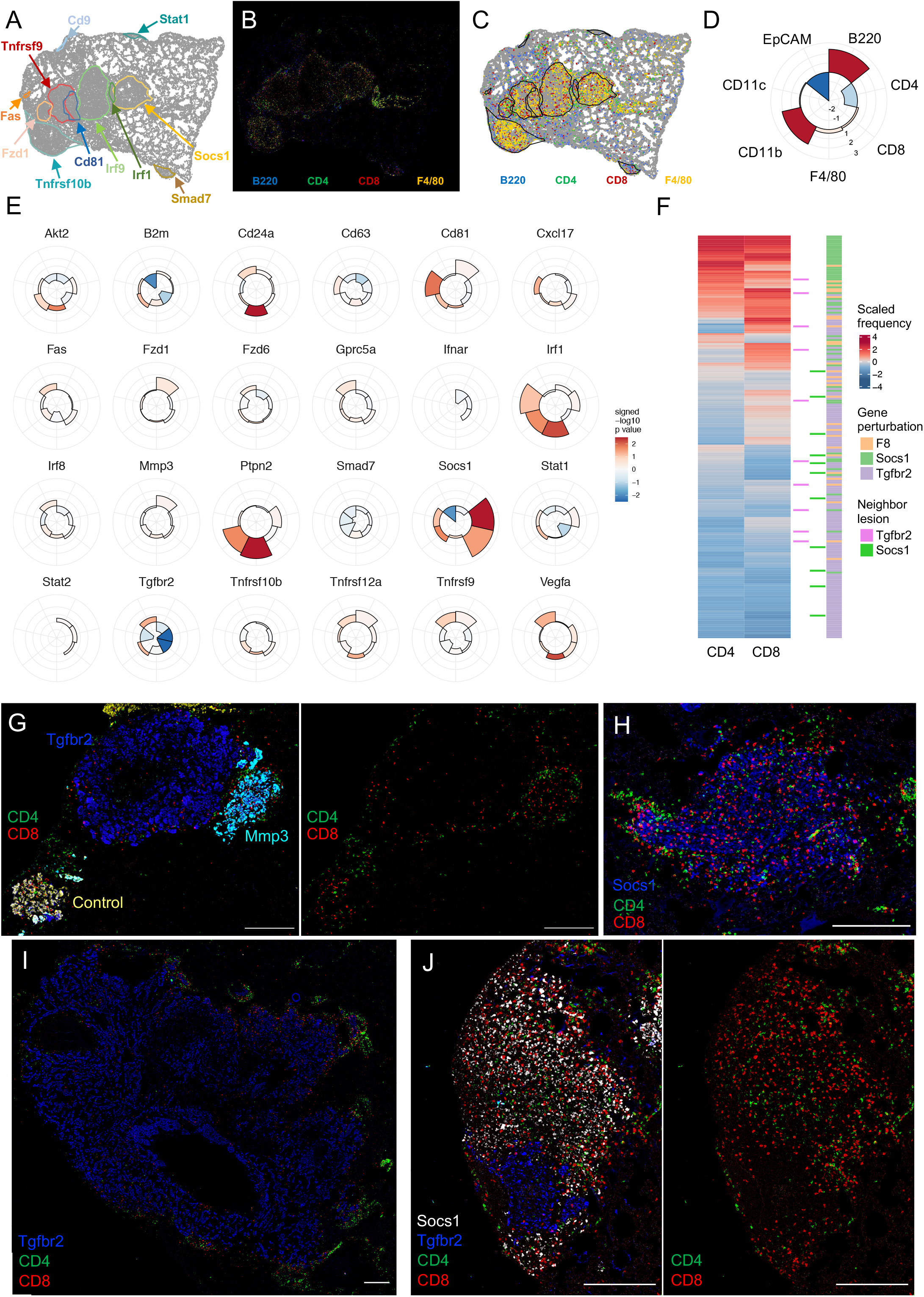
Perturb-map correlates gene perturbation with archetypes of immune infiltration. (**A-D**) Overview of the Perturb-map analysis pipeline for the quantification of immune infiltration and EpCAM expression. Analysis was performed on lung tissue collected from mice described in Figure 3 (n=11 mice, 2 separate experiments). (**A**) Representative example of an in silico reconstituted image displaying tumor boundaries (same section as Figure 3C-E, digital image shown is from the right panel of Figure 3D for perspective). (**B**) Representative image of immune infiltration in KP lung tumor lesions. Lung tissue sections that were stained the Pro-Code epitopes were stained for B220, CD4, CD8, CD11b, CD11c, F4/80 and EpCAM using MICSSS. The image represents a pseudo-colored overlay of B220, CD4, CD8 and F4/80, color-coded as indicated. (**C**) Cells positive for each immune markers are displayed in the in silico Pro-Code tumor map. Cell densities per mm^2^ were subsequently quantified within each tumor lesion. (**D**) Schematic of a radial plot used to visualize the immune landscape associated with each gene perturbation. Each fraction corresponds to immune cell density, or mean intensity of EpCAM staining, within the tumor border (as visualized in panel C). Bar height represents the difference in median cell density or mean intensity between control tumors (F8, n=55) and tumors carrying gene perturbations, rescaled as a Z-score for each marker. Opacity of the shading represents the −log10 adjusted p value of a Wilcoxon rank-sum test of the same comparison signed by the direction of effect. (**E**) Analysis of immune infiltration and exclusion in Pro-Code/CRISPR lung tumors. Shown are radial plots indicating immune density (and mean EpCam expression) associated with tumors with the indicated gene perturbation relative to control KP tumors (F8, n=55). Shown are gene perturbations with >20 lesions. (**F**) Examination of T cell density in F8, Socs1 or Tgfbr2 tumors related to their proximity to Socs1 or Tgfbr2 tumors. The heatmap represents the Z-score of log transformed CD4^+^ and CD8^+^ T cells density (censored at the 95^th^ percentile) in control (F8), Socs1 and Tgfbr2 tumors. Tumors neighbored by a Socs1 or Tgfbr2 lesion (boundary distance inferior to 75 mm) are identified. (**G-J**) Representative examples of CD4^+^ (green) and CD8^+^ (red) T cell infiltration within indicated tumor lesions. Annotated gene perturbations were identified based on Pro-Code expression. Scale bars, 100 μm.

Specific patterns of immune infiltration were found in lesions with different gene perturbations (**Figure 4E**). For example, in tumors in which the CRISPR targeted B2m, which is required for antigen-dependent interactions with CD8^+^ T cells, there was a specific reduction in CD8^+^ T cells, supporting the need for TCR/pMHC interactions to maintain T cells in the TME. Loss of Irf1 and Ptpn2 both led to increased infiltration of myeloid cells (F4/80+, CD11b+), and to a lesser extent CD4+ T cells. Strikingly, tumors in which Tgfbr2 or Socs1 were targeted, which were both enriched in vivo, displayed inverted patterns of immune composition. Tgfbr2 lesions were markedly excluded of immune cells, especially CD4+ and CD8+ T cells (**Figure 4E, 4F, 4G, 4I and 4J**), whereas Socs1 lesions were highly infiltrated by T cells (**Figure 4E, 4F, 4H and 4J**). Socs1 tumors also had lower levels of EpCAM expression. This is suggestive of altered differentiation, which is consistent with the histopathology of Socs1 tumors that were predominately found to be moderate to poorly differentiated (**Figure S3B and S3C**).

Metastatic lesions can evolve divergent molecular states and even within a single tumor mass genetic sub-clones can form spatially distinct regions with different TME composition (Mitra et al., 2020), but how adjacent, genetically heterogeneous regions might influence each other is not well explored. We sought to use Perturb-map to investigate whether neighboring tumors influenced the immune infiltration of one another. We identified control (F8), Socs1 and Tgfbr2 tumors that were in contact with a neighboring Tgfbr2 or Socs1 tumor and clustered them based on T cell infiltration. Socs1 tumors were dominant in lesions highly infiltrated by T cells, whereas Tgfbr2 tumors were mainly excluded in T cells (**Figure 4F and 4G, 4H and 4I**), as we had observed from our previous analysis (**Figure 4E**). Notably, tumor lesions with an adjacent Socs1 KO tumor did not tend to display higher T cell infiltration, while tumor lesions neighbored by Tgfbr2 KO tumors were not more excluded in T cells. Even when Tgfbr2 KO lesions were directly adjacent to a highly infiltrated Socs1 KO lesion, the Tgfbr2 KO lesion remained immune excluded (**Figure 4J**). This suggests, at least in the case of IFNγ and TGFβ pathway alterations, that in lung tumor lesions in close contact with each other, the composition and spatial arrangement of immune cells is shaped very locally. Though this is likely dependent on the specific molecular alterations, as some gene expression changes may have more distal effects, Perturb-map can provide a scaled means to assess how specific genetic alterations influence local, proximal, and distal TME states.

### Perturb-map paired with spatial transcriptomics identifies perturbation associated molecular signatures

To further determine the influence of different gene perturbations on tumor state, we paired spatial transcriptomics with Perturb-map. We applied the 10X Visium technology on four sections of mouse lungs seeded with KP tumors expressing the Pro-Code/CRISPR library, as described above (**Figure 3B**). Differential expression of kmeans derived clusters distinguished a specific gene signature in spatial domains corresponding to tumor lesions (**Figure 5A and Figure S4A**). In contrast to healthy surrounding lung tissue, tumor lesions highly expressed specific keratins and epithelial markers (e.g. Krt8, Krt18, Epcam) (**Figure S4B**). We also found evidence of an IFNγ signaling signature, including upregulation of the antigen presentation pathway (e.g. Nlrc5, Tap1, Psmb8, Psmb9), IFNγ signaling pathway genes (e.g. Stat1, Irf1, Socs1), and IFNγ-induced chemokines (e.g. Cxcl9, Cxcl10) (**Figure S4B)**. Of note, Pro-Code transcripts were captured distinctly and specifically localized within tumor cluster regions (**Figure 5B**).

**Figure 5.**
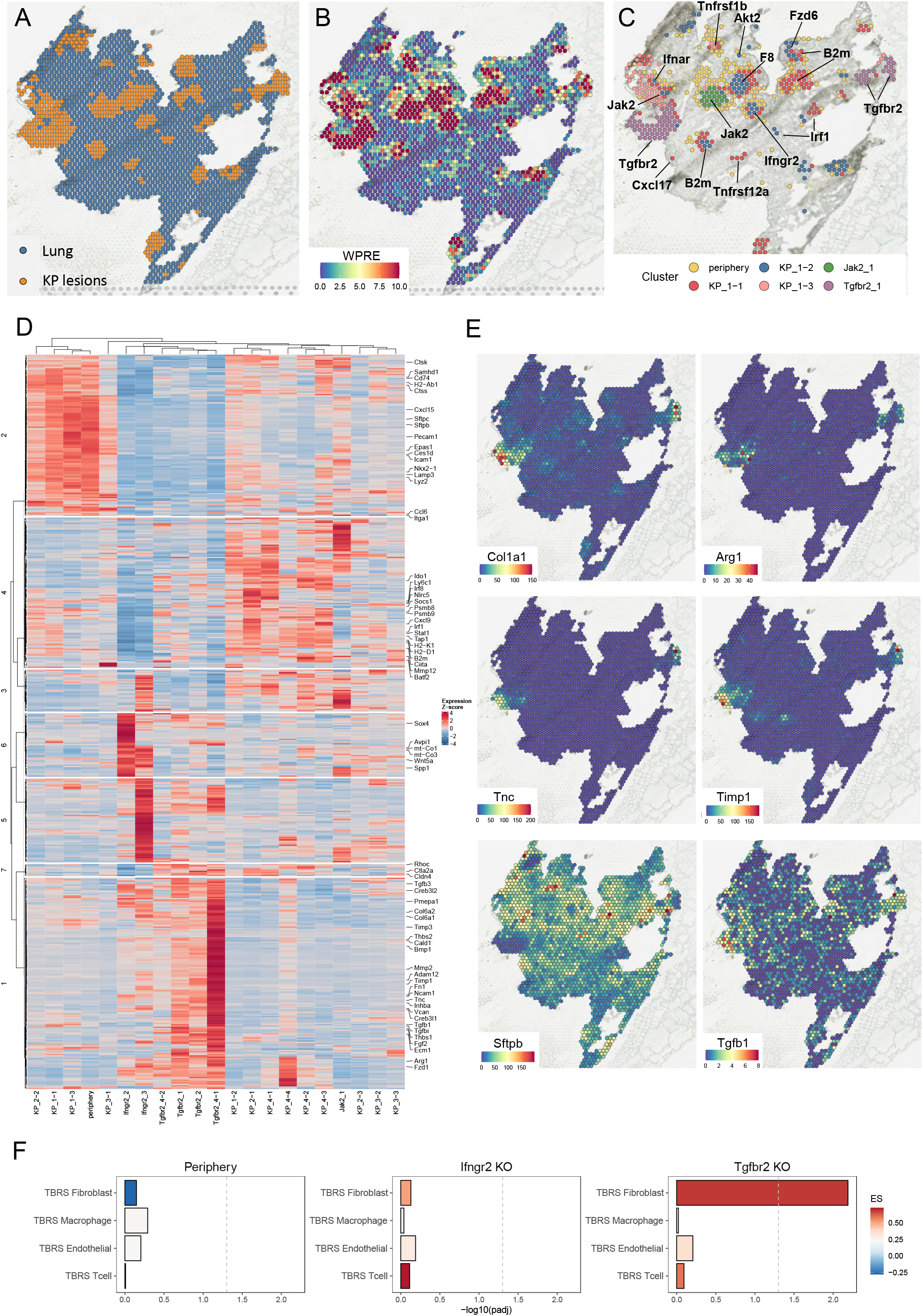
Perturb-map with spatial transcriptomics identifies perturbation-specific gene signatures. (**A**) Kmeans clustering (k = 2) of KP lesions and non-involved lung expression spots from a representative tissue section profiled by spatial transcriptomics (10X Visium). The same representative tissue section is used for panels A-C. (**B**) Unique Molecular Identifier (UMI) counts of reads mapping to the WPRE region of the Pro-Code transcript in each Visium spot. (**C**) Leiden clustering of Pro-Code+ spots (>= 4 WPRE UMIs). Gene perturbations were annotated based on imaging mass cytometry staining of a serial section. (**D**) Gene expression modules derived from differentially expressed genes (adjusted p value < 0.01 and log_2_ fold-change > 0.5) in Ifngr2 and Tgfbr2 spot clusters relative to clusters representing the KP lesion signature. Genes and Leiden spot cluster means were arranged by hierarchical clustering using Pearson correlation distance and average agglomeration. Gene clustering was performed with all Leiden spot clusters except Jak2_1 and KP_3-1 and a tree cut with 7 groups was used to define modules. The color corresponds to the row Z-score of mean log-transformed SCTransform corrected counts for the indicated Leiden spot clusters. These values were used for hierarchical clustering of genes, whereas the mean scaled Pearson residuals for each feature in each Leiden cluster was used for clustering of tumor groups. (**E**) UMI counts for select transcripts differentially expressed in the Tgfbr2 spots. (**F**) −log_10_ adjusted p value of a gene set enrichment analysis in TGFβ response score (TBRS) categories defined in Calon et al. Average log_2_ fold-change of differentially expressed genes (Bonferroni adjusted p-value < 0.01) in the listed signature were used as a test input. Enrichment Score (ES) indicates the magnitude and direction of effect for the enrichment.

Graph-based clustering of tumor spots defined by Pro-Code transcripts revealed distinct and repeated tumor gene expression signatures across lung sections (**Figure 5C and Figure S4C**). Consistent with the clonal development of KP tumors, spatial domains corresponding to a given lesion often clustered together, but there were also specific lesions that clustered distinctly from the rest. To identify the gene perturbations in each lesion, we imaged a serial section of lung tissue stained with metal-conjugated antibodies to each Pro-Code epitope tag by imaging mass cytometry (Hyperion) (**Figure S4D**). Similar to MICSSS and MIBI, Hyperion was able to resolve each epitope tag with a high signal to noise ratio. The identification of Pro-Code/CRISPRs within the tissue allowed us to annotate each gene signature with the corresponding gene perturbation, and revealed that many tumors carrying CRISPRs targeting Tgfbr2 or Ifngr2 presented a distinct gene signature compared to other KP tumors (**Figure 5C and Figure S4C**).

Hierarchical clustering of tumor cluster mean expression values across Visium sections, which contained tumor-bearing lung tissues from separately injected mice, indicated that the gene signatures associated with particular perturbations or tissue regions were highly consistent (**Figure 5D**). Additionally, clustering of the genes themselves that were significantly changing in Ifngr2 and Tgfbr2 KO tumor clusters highlighted patterns consistent with orthogonal phenotyping from our image-based analysis (**Figure 5D**). Spatial domains located at the periphery of tumor lesions presented a distinct inflammatory signature, characterized by the upregulation of Cxcl15, a neutrophil attractant, Lyz2 and Lamp3, markers of myeloid cells, and high levels of the MHC Class II machinery, including H2-Ab1 and Cd74. Ifngr2 KO tumors exhibited a broad down-regulation of genes involved in antigen presentation (H2-k1, H2-d1, Ciita, Nlrc5, Tap1, Psmb8, Psmb9) as well as known IFNγ-induced genes, such as Irf1 and Cxcl9.

In Tgfbr2 KO tumors there was an increase of various collagen-coding genes, including Col1a1, Col6a1, and Col6a2 (**Figure 5D and 5E**), consistent with the presence of a fibro-mucinous stroma (**Figure 3H and Figure 3I**). Unexpectedly, many of the genes upregulated in the Tgfbr2-targeted tumors were indicative of increased TGFβ signaling, including Tnc, Timp1, and Arg1, which are induced by TGFβ (Calon et al., 2012; Kwak et al., 2006; Mondanelli et al., 2017; Yoon et al., 2021) (**Figure 5E**). There was also downregulation of Sftpb, which is negatively regulated by TGFβ (Wesselkamper et al., 2005). Notably, Tgfb1 and Tgfb3, ligands of Tgfbr2, were also upregulated in the Tgfbr2 tumors lesions (**Figure 5D and 5E**).

The Visium platform has a spot diameter of 55μm which means that the transcriptomic profile of each capture region is derived from several cells, and these can be of distinct type, including cancer, immune, endothelial, and stroma. To better understand which cell types might be responding to TGFβ in the Tgfbr2 KO tumors, we compared the differentially expressed genes in these lesions with a published dataset that reported the TGFβ signatures of fibroblasts, endothelial cells, macrophages and T cells (Calon et al., 2012). While there was relatively little overlap between the genes upregulated in the Tgfbr2 KO tumors and TGFβ treated macrophages and T cells (consistent with the dearth of these cells in the tumor), there was a significant overlap with TGFβ treated fibroblasts (**Figure 5F**), suggesting increased TGFβ signaling was occurring in the stroma of Tgfbr2 KO tumors.

These findings demonstrate that a consequence of Tgfbr2 loss on KP lung cancer cells is increased TGFβ ligand expression in the tumor along with TGFβ pathway activation in the TME, and suggests a potential mechanism for the stroma remodeling and immune exclusion we observed in the Tgfbr2 KO lesions. They also demonstrate that spatial transcriptomics can be incorporated into Perturb-map to provide unbiased molecular analysis of different gene perturbations in a spatially resolved manner.

### Validation that loss of Tgfbr2 results in more aggressive and T cell excluded KP lung tumors

We sought to confirm the Perturb-map phenotype associated with Tgfbr2 targeted KP tumors. To do this, we transduced KP-Cas9 cells with LV encoding CRISPR gRNAs targeting either F8 (control) or Tgfbr2, and injected the cells i.v. into Cas9 mice. We harvested lungs at 7, 14, and 28 days after injection and examined tumor burden. Within 7 days of injection there was already evidence of larger and more abundant Tgfbr2 KO tumors compared to control. This was even more apparent at later time points in which individual Tgfbr2 KO tumor lesions were >2-fold bigger than control lesions, resulting in ∼10% of lung area being covered by Tgfbr2 tumors compared to ∼4% for the control (**Figure 6A**).

**Figure 6.**
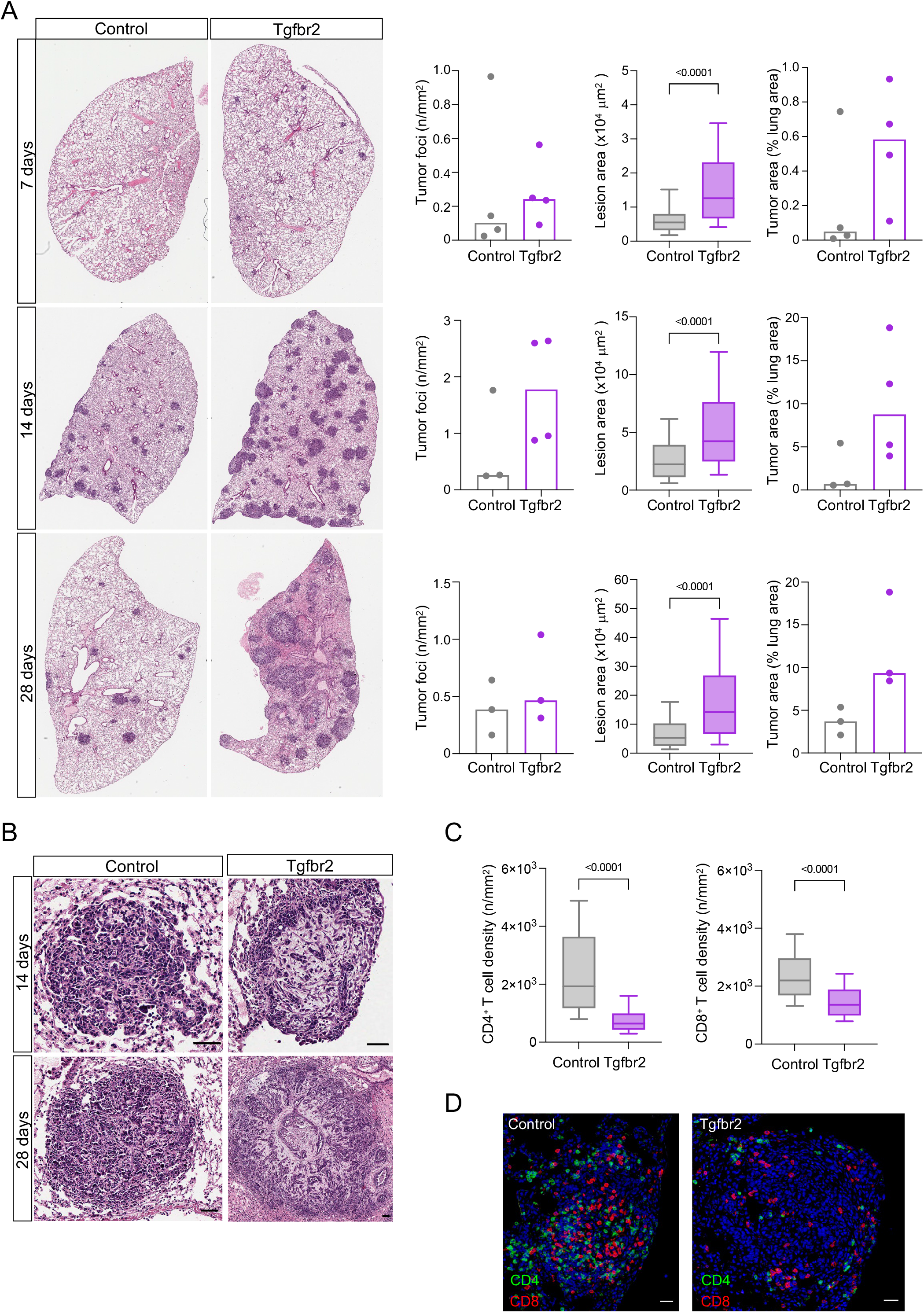
Kinetics of Tgfbr2 KO and control KP lung tumor development. (**A**) Comparison of lung tumor burden between mice bearing control or Tgfbr2 CRISPR KP tumors. KP-Cas9 were transduced with vectors encoding CRISPRs targeting F8 (control) or Tgfbr2 and subsequently injected i.v. into Cas9 mice (n=3-4 mice/group/time point). Lungs were collected at 7, 14, and 28 days. Whole lung sections were stained with H&E. A representative lung lobe is shown for each condition. Density of tumor lesion (left graph), lesion size (middle graph) and total tumor area (right graph) were quantified. Statistical significance was determined using a Mann-Whitney test. (**B**) Representative examples of control or Tgfbr2 targeted tumor lesions. Scale bar, 50 mm. (**C**) Quantification of CD4^+^ (left) and CD8^+^ (right) densities in tumor lesions across all samples at the 14-day time point. Statistical significance was determined using a Mann-Whitney test. (**D**) Representative image of T cell infiltration in a control or Tgfbr2 targeted tumor lesion. Scale bar, 50 mm.

Similar to what we observed with Perturb-map, many of the Tgfbr2 tumors had a fibro-mucinous stroma (apparent in some lesions by the 2-week time point), which was not observed with control tumors (**Figure 6B**). In addition, as we had observed in the Perturb-map screen, there was a significant decrease in CD4+ and CD8+ T cell infiltration within Tgfbr2 tumors (**Figure 6C and 6D**). These studies provide an indication that Perturb-map can be used to reliably identify complex tumor phenotypes induced by gene perturbations at scale.

## DISCUSSION

These studies establish a new approach for functional genomics which enables pooled CRISPR screens to be resolved within tissue by multiplex imaging and spatial transcriptomics. A key advance of Perturb-map is that it broadens the types of genes and gene functions that can be investigated in functional genomics screens. Most sequencing-based pooled screening approaches require dissociation of cells or tissues for analysis, and this results in loss of information about how a perturbation might affect a cell’s local environment. Indeed, for standard CRISPR screens, identifying the CRISPRs requires cell destruction to collect DNA for barcode analysis, and this prevents complex cell phenotyping as a readout. As noted above, these caveats have largely restricted CRISPR genomics to screening for cell intrinsic gene functions, often related to how a gene influences cell fitness under a selective pressure or expression of a specific phenotypic marker (Doench, 2018; Shalem et al., 2015). By imaging the Pro-Codes in tissue, Perturb-map enabled us to assess how different gene perturbations not only altered tumor fitness, but also tumor morphology and immune cell recruitment. To our knowledge, this is the first CRISPR genomics platform that has demonstrated the capability for this type of phenotypic screening.

Our initial application of Perturb-map was intended to identify genes affecting immune cell recruitment. We focused on a small set of immune cell types for analysis, but any cell type or protein that can be detected by antibody staining could be included for study. As there are multiplex imaging techniques and instruments that can detect as many as 60 proteins on a single tissue section, much more extensive detection of cell types and cell states is possible (Goltsev et al., 2018; Jackson et al., 2020; Lin et al., 2018). It is worth noting that Perturb-map, as presented here using MICSSS, does not require any special instrumentation or reagents and should therefore be feasible for most labs to carry out (Akturk et al., 2020). Even the antibodies used to detect the Pro-Code are widely available, as the majority of epitopes are commonly used tags, such as FLAG, HA, StrepII, and AU1 (Wroblewska et al., 2018).

Beyond examining tumor morphology and immune cell recruitment, the ability to readout CRISPR screens by imaging can enable scaled studies to identify genes that control many other processes important to tumor biology, such as protein localization in a cell (e.g. cytoplasmic vs nuclear), tumor organization (e.g. cell proximity to vasculature or stroma), cell migration and invasion (e.g. to determine cancer cell invasion in to normal tissue), and cell-cell interactions (e.g. to identify molecules mediating contact between cells). Additionally, because Perturb-map is compatible with tissue analysis, it can be applied to study genes dependent on factors only available in vivo. For example, a cell receptor may have little function in vitro because its ligands are not present in the culture media but will mediate signaling in a tissue in which stromal cells express its ligand.

The coupling of spatial transcriptomics with Perturb-map provided an unbiased means to determine how specific gene perturbations altered the molecular state of a tumor. This expands the possible applications of Perturb-map for discovering gene functions and can help provide mechanistic insights into phenotypic observations, as we found in Tgfbr2 targeted lesions. The scale relative to cost of current commercial spatial transcriptomics platforms is a limitation for analyzing many perturbations, as the region of analysis is relatively small (6.5mm^2^ per Visium slide). For perspective, whole slide scanning for techniques such as MICSSS can image >1500mm^2^ of tissue for a nominal cost. Future advances in spatial transcriptomic technology will address these limitations including improved cellular resolution (Cleary et al., 2021; Larsson et al., 2021), but a multiscale screening approach may still be optimal, as we used here: employing multiplex imaging to screen for defined phenotypes of interest (e.g. altered immune cell recruitment, hypoxia, tissue invasion, etc.) on larger and more sections and spatial transcriptomics for more comprehensive molecular analysis on limited tissue regions.

We also demonstrated that we could resolve memPC and nPC in the same cell. This provides the possibility of a scaled means to assess genetic interactions. Cells can be transduced with two different libraries to generate random multi-gene KOs, which could be resolved in vivo with spatial resolution. Even just combining two different Pro-Code/CRISPR libraries targeting 35 different genes would generate 1,225 double gene perturbations. This can be valuable for identifying redundant gene pairs and cooperating gene functions (Horlbeck et al., 2018; Mair et al., 2019). A challenge of this type of analysis with single cell approaches is sampling enough events. However, this should be feasible with modest sized libraries using Perturb-map because of the high number of cells in a tissue that can be imaged by whole slide scanning. In addition to functional genomics, the ability to combine different Pro-Code libraries provides considerable scale for spatial mapping of cancer clones in situ. The Pro-Codes could be used to genetically label 100s of different cancer lines or patient-derived cancer cells, and study their growth, biology, and competition in vivo, similar to what has been done with DNA barcodes (Jin et al., 2020; Wagenblast et al., 2015), but with the ability to resolve different clones spatially.

We used Perturb-map to screen the contributions of specific genes on lung cancer growth. The de-enrichment of Pd-l1 targeting Pro-Code/CRISPR tumors supported the utility of the approach for investigating tumor immunity. Interestingly, Cd47 targeted tumors were also significantly de-enriched, suggesting KP tumors protect themselves from macrophage clearance with the CD47 ‘don’t eat me signal’. We recently reported that tissue-resident macrophages provide a supportive niche during early lung tumor growth (Casanova-Acebes et al., 2021), and it may be that CD47 is needed to keep this beneficial relationship from becoming carnivorous.

We also found that suppression of IFNγ signaling provided KP tumors a growth advantage. This was somewhat unexpected as there is good evidence that IFNγ is important for tumor immunity, and mutations in JAK/STAT have been associated with resistance to checkpoint blockade (Gao et al., 2016; Zaretsky et al., 2016). However, a more complicated view of IFNγ’s role has emerged which indicates it can have both pro and anti-tumor effects (Benci et al., 2019; Gocher et al., 2021). The opposing effects are believed to be related to IFNγ’s activity in inducing both antigen presentation pathways and checkpoint ligands in cancer cells, as well as IFNγ’s different effects on cancer and immune cells. Whereas most in vivo studies have examined the consequence of loss of the pathway (e.g. through Ifngr2 or JAK/STAT knockout), our results indicate that enhancement of IFN γ signaling specifically in cancer cells, through Socs1 knockout, can facilitate tumor growth. Notably, increased growth occurred despite increased T cell infiltration in the Socs1 lesions. As Socs1 loss leads to upregulation of PD-L1(Wroblewska et al., 2018), it is likely that the Socs1 cancer cells were able to survive by promoting T cell exhaustion. If this is the mechanism, Socs1 could be a good therapeutic target to pair with PD1/PDL1 blockade, as SOCS1 inhibition may enhance T cell infiltration while the effects of upregulated PD-L1 can be blunted.

Our Perturb-map analysis found that Tgfbr2 targeted tumors were more abundant, larger, and mucinous than control tumors, and had excluded T cells, which we confirmed occurred even at early stages. While TGFBR2 mutations are not prevalent in lung cancer, several studies have shown that TGFBR2 mRNA and protein are markedly downregulated in human lung tumors, and downregulation correlates with poor survival (Borczuk et al., 2005; Malkoski et al., 2012). Different functions of TGFBR2 on lung cancer cells likely contribute to this outcome, but our data suggest that immune exclusion may play a role. TGFβ has been shown to promote immune exclusion and TGFβ blockade is a promising cancer immunotherapy strategy which is currently being evaluated in clinical trials (Mariathasan et al., 2018). Thus, it was counterintuitive that loss of the receptor resulted in cold tumor lesions. However, the spatial transcriptomics seemed to reconcile these findings by indicating that TGFβ and TGFβ signaling was upregulated in the TGFβ KO lesions. Thus, despite the cancer cells not being responsive to TGFβ, the immune and stroma cells were still TGFβ sensitive and subject to its suppressive and remodeling effects (Derynck et al., 2021). In addition to TGFβ being upregulated, loss of the TGFβ receptor on the dominant cell type in the lesion, the cancer cells, may have had the effect of increasing the bioavailable TGFβ in the lesion, further propagating TGFβ’s effect on the immune and stroma compartments. Thus, one of the consequences of downregulation of TGFBR2 in patient lung tumors (Borczuk et al., 2005; Malkoski et al., 2012) may be increased immune suppression.

Beyond investigating specific gene functions, Perturb-map also enabled us to examine how specific gene perturbations and associated phenotypes influenced neighboring tumor regions. This is a relatively underexplored area of tumor biology, but as more high-resolution studies report heterogeneity in the TME and associations with tumor genetics (Lomakin et al., 2021; Mitra et al., 2020), Perturb-map can provide a valuable means for causal studies to determine how particular genes influence local and distal regions of a tumor.

## ACKNOWLEDGMENTS

We thank R. Samstein and J. Aguirre-Ghiso (Mount Sinai) for helpful discussions. We also thank P. Suri, D. D’souza and R. Sweeney (Mount Sinai), the Mount Sinai Human Immune Monitoring Center (HIMC), and O. Elemento and H. Ravichandran (Englander Institute of Precision Medicine Mass Cytometry Core) for technical assistance. B.D.B. was supported by NIH (R33CA223947 and R01AT011326) and the Cancer Research Institute. M.M. was supported by NIH (R01CA257195 and R01CA254104). M.D. was supported by the Belgian American Educational Foundation and S.A.R. was supported by NIH (T32AI007605), the Gladys and Roland Harriman foundation and the Wrobel Family Foundation.

## AUTHOR CONTRIBUTIONS

Conceptualization, M.D. and B.D.B.; Methodology, M.D., S.A.R., G.A., A.W.; Software, S.A.R., M.B.; Investigation, M.D., A.W., E.S.P., S.R.N., L.P., J.L.B., A. Bektesevic, B.H.L.; Writing – Original Draft, M.D. and B.D.B.; Writing – Review & Editing, M.D., S.A.R., M.M., B.D.B.; Supervision, A. Baccarini, N.B., A.H.R., S.G., D.P., M.M.; Funding Acquisition: M.M. and B.D.B.

## MATERIAL AND METHODS

### EXPERIMENTAL MODEL AND SUBJECTS DETAILS

#### Mice

BALB/cJ (stock number 000651), C57BL/6J (stock number 000664) and Cas9 (stock number 027650) mice were purchased from Jackson Laboratory. At the time of experimentation, mice were 8-12 weeks of age. All mice were hosted in a specific pathogen-free facility. The Institutional Animal Care and Use Committee of the Icahn School of Medicine at Mount Sinai reviewed and approved all animal protocols used in the present study.

#### Cell culture

293T cells (embryonic kidney; human) and KP cells (lung adenocarcinoma; C57BL/6 mice) (Xue et al., 2011) were grown in IMDM with 10% heat-inactivated FBS (GIBCO), 100 U/ml penicillin/streptomycin (GIBCO) and 2 mM L-Glutamin. 4T1 cells (mammary carcinoma; BALB/c mice) were grown in RPMI with 10% heat-inactivated FBS (GIBCO), 100 U/ml penicillin/streptomycin (GIBCO) and 2 mM L-Glutamin. Each passage, cells were washed with PBS, detached from the plate with 0.05% Trypsin-EDTA (GIBCO) and replated. Cells were discarded after 12 passages. 293T and 4T1 were purchased from ATCC. KP cells stably expressing Cas9 (KP-Cas9) were engineered by transduction with Cas9 lentivirus as previously described (Wroblewska et al., 2018).

### METHOD DETAILS

#### Vector construction

The nuclear Pro-Codes (nPC) vectors were cloned following the same structure as the NGFR Pro-Codes (Wroblewska et al., 2018). The NLS from hnRNPK (21-KRPAEDMEEEQAFKRSR-37) (Matunis et al., 1992) was fused to the 3’ end of the coding sequence for mCherry by PCR. The mCherry-NLS was cloned downstream of the EF1a promoter into a lentiviral vector that contained a U6 gRNA expression cassette. The linear epitope combination sequences were cloned at the 5’ end of the mCherry sequence using BamHI and SphI restriction sites. To clone gRNA sequences, Pro-Code vectors were digested with BbsI, purified using PCR purification kit (QIAGEN) and ligated with pairs of annealed oligo sequences as described previously (Wroblewska et al., 2018). gRNA sequences were obtained from the Brie CRISPR library (Doench et al., 2016). TOP10 competent cells were used for all subsequent plasmid preparations. All plasmids were purified using ZR Plasmid Miniprep Classic kit (Zymo Research) or EndoFree Plasmid Midi Kit (QIAGEN).

#### Pro-Code/CRISPR libraries

The following genes were targeted in the Pro-Code/CRISPR library used in Figure 2 to 4: Akt1, Akt2, B2m, Cd24a, Cd47, Cd63, Cd81, Cd9, Cxcl17, F8, Fas, Fzd1, Fzd6, Gprc5a, Ifnar, Ifngr2, Irf1, Irf8, Irf9, Jak1, Jak2, Mmp3, Pd-l1, Ptpn2, Smad7, Socs1, Stat1, Stat2, Tgfbr2, Tnfrsf10b, Tnfrsf12a, Tnfrsf1a, Tnfrsf1b, Tnfrsf9, Vegfa. For each targeted gene, 3 plasmids were generated, corresponding to 3 different gRNA cloned into the same Pro-Code backbone.

#### Lentiviral vector production and titration

Lentiviral vectors were produced as previously described in detail (Baccarini et al., 2011). For Pro-Code/CRISPR libraries, lentiviral vectors production was arrayed in 6-well plates. 293T cells were seeded at 500,000 cells per well 24 hours before calcium phosphate transfection with third-generation VSV-pseudotyped packaging plasmids and the transfer plasmids. Each well received 3 transfer plasmids in equimolar amount, encoding 3 different gRNAs targeting the same gene and coupled to the same Pro-Code. Supernatants were collected 48 hours post-transfection, strained through a 0.22-µm PVDF disc filter and stored at −80°C.

#### Vector transduction

4T1 and KP cells were transduced as previously described (Wroblewska et al., 2018). Briefly, cells were seeded 24 hours prior to transduction at a density of 50,000 cells per well in 6-well plates and transduced with lentiviral vectors in the presence of 5 µg/ml polybrene (Millipore). When cells were transduced with a Pro-Code library, cells were transduced at a low MOI to ensure that a majority of cells received only one vector (< 10% Pro-Code positive cells). Alternatively, KP-Cas9 cells were transduced in array with each Pro-Code/CRISPR lentiviral vector mix (3 plasmids, each coding for a gRNA targeting the same gene) and transduced at 60 MOI to achieve 60-70% Pro-Code positive cells which were subsequently sorted to >99% purity (below).

#### Flow and Mass cytometry

Adherent cells were detached with 0.05% trypsin-EDTA and resuspended in PBS. For analysis of transduction efficiency and sorting of Pro-Code positive cells, cells were stained with anti-human CD271 PE or Alexa Fluor 647 (BD Biosciences) and flow sorted based on NGFR or mCherry expression. The KP-Cas9 cells transduced with Pro-Code/CRISPR lentiviral vectors and used in Figures 2 to 4 were sorted into the same tube (1 x10^4^ cells/Pro-Code).

Processing and analysis of cell suspension by mass cytometry was performed as described previously in detail (Wroblewska et al., 2018). Briefly, cell suspensions were stained for viability with Cell-ID Intercalator-103 Rh for 15 min at 37°C. Cells were subsequently stained for surface markers in flow buffer supplemented with anti-mouse CD16/CD32 antibody (eBioscience) for 30 min on ice. Next, cells were fixed and permeabilized using either BD Cytofix/Cytoperm solution (BD Biosciences) or the eBiosciences Intracellular Fixation & Permeabilization kit (Thermo Fisher Scientific) and stained with epitope-tag specific antibodies for 30 min on ice. Cells were then washed and incubated in 125 nM Ir intercalator (Fluidigm) diluted in PBS containing 2.4% formaldehyde for 30 min at RT, washed and stored in PBS at 4°C until acquisition. The samples were acquired on either a CyTOF2 or Helios (both Fluidigm) at an event rate of < 500 events/second. The following antibodies were used for CyTOF staining: anti-HA tag-147Sm (clone 6E2, Cell Signaling), anti-V5 tag-152 Sm (Thermo Fisher Scientific), anti-DYKDDDDK (FLAG) tag-175Lu (Clone L5, Biolegend), anti-VSVg tag-158 Gd (rabbit pAb, Thermo Fisher Scientific), anti-E tag-154Sm (clone 10B11, Abcam), anti-NWSHPQFEK (NWS) tag-159Tb (clone 5A9F9, Genscript), anti-AU1-162Dy (clone AU1, BioLegend), anti-AU5-169Tm (clone AU5, BioLegend), anti-Ollas tag-153Eu (clone L2, Thermo Fisher Scientific), anti-HSV tag-172Yb (rabbit polyclonal, Thermo Fisher Scientific), anti-S tag-165Ho (clone SBSTAGa, Abcam), anti-Protein C tag-171Yb (clone HPC4, Genscript), anti-mCherry-142Nd (Abcam). Antibodies were purchased purified and conjugated in-house using MaxPar X8 Polymer Kits (Fluidigm) according to the manufacturer’s instructions.

#### Tumor models

4T1 murine mammary gland carcinoma cells were injected (5×10^4^ cells) in the mammary fat pad of 8-12 week old BALB/c WT mice. Mice were sacrificed 12 days post-inoculation and the primary tumor was excised. Alternatively, mice were sacrificed after 28 days and the lungs were collected. KP cells were injected (1×10^6^ cells) intravenously (i.v.) in the tail vein of C57BL/6 mice or Cas9 mice (to avoid immune rejection of Cas9 expressing KP cells). Mice were sacrificed 28 days later, and the lungs were collected. All the collected samples were either fresh-frozen in OCT or fixed overnight in PLP buffer (0.075 M lysine, 0.37 M sodium phosphate, 2% formaldehyde and 0.01 M NaIO_4_, at pH 7.2) and paraffin-embedded.

#### Pathological Assessment of tumor lesions

Paraffin-embedded lung sections were stained with hematoxylin/eosin and scanned using a Aperio AT2 slide scanner (Leica). The gRNA present in each tumor lesion was identified based on the Pro-Code expression. Each tumor lesion was delimited in QuPath and evaluated by a pathologist (blinded to the tumor identification). The assessed criteria were tumor location, degree of differentiation and stromal composition. Specific patterns of tumor development were also characterized.

#### Imaging Mass Cytometry

Sections from fresh frozen tissue were fixed in 2% paraformaldehyde for 30 minutes on ice, blocked with 5% normal goat serum in TBS 2% Tween 20 (TBS-T) at RT for 15 min and incubated overnight at 4°C with metal-conjugated antibodies. Sections were then washed with TBS-T and fixed in 2% glutaraldehyde for 5 minutes. Following 3 washes in Tris buffer (pH 8.5) and deionized water, sections were dehydrated in ascending series of 50%, 75%, 95% and 100% ethanol for 1 minute each. Slides were then dried at RT and stored into a vacuum chamber until acquisition. The following antibodies were used: anti-HA tag-147Sm (clone 6E2, Cell Signaling), anti-V5 tag-152 Sm (Thermo Fisher Scientific), anti-DYKDDDDK (FLAG) tag-175Lu (Clone L5, Biolegend), anti-VSVg tag-158 Gd (rabbit pAb, Thermo Fisher Scientific), anti-E tag-154Sm (clone 10B11, Abcam), anti-NWSHPQFEK (NWS) tag-159Tb (clone 5A9F9, Genscript), anti-AU1-162Dy (clone AU1, BioLegend), anti-AU5-169Tm (clone AU5, BioLegend), anti-Ollas tag-153Eu (clone L2, Thermo Fisher Scientific), anti-HSV tag-172Yb (rabbit polyclonal, Thermo Fisher Scientific), anti-S tag-165Ho (clone SBSTAGa, Abcam), anti-Protein C tag-171Yb (clone HPC4, Genscript), anti-mCherry-142Nd (Abcam). Samples were acquired with the Hyperion Imaging System (Fluidigm) or a MIBIscope System (Ionpath). Data related to Figure 1 (and associated supplementary figures) was acquired on a MIBIscope system (Ionpath). Tissue sections were cut onto a gold-plated slide (Ionpath). Fields of view of 500 um^2^ were acquired using the optimal resolution setting with a 1024 x 1024 pixel image resulting in a 488 nm resolution. The ion beam dwell time for each field of view was 3 ms per pixel with 4 repeat scans for a total dwell time of 12 ms per pixel. Images were then processed using the IONpath MIBI/O software to generate TIFF images which were further cleaned for isobaric correction and filtered to substract background noise signal. Images related to Figure 5 (and associated supplementary figure) were acquired on the Hyperion Imaging System (Fluidigm). Region of interests for laser ablation were selected based on the location of Pro-Code positive tumors identified on a serial section by spatial transcriptomics (Visium, 10x Genomics). All ablations were performed with a laser frequency of 200 Hz. The raw MCD files were exported for analysis on the MCD Viewer Software (Fluidigm).

#### Multiplexed Immunohistochemical Consecutive Staining on Single Slide (MICSSS)

Iterative cycles of immunostaining on 4-µm thick FFPE tissue sections were performed as described previously in detail (Akturk et al., 2020). Briefly, slides were baked overnight at 60°C, then de-paraffinized in xylene and rehydrated in descending series of 100%, 90%, 70% and 50% ethanol. Slides were incubated at 95°C for 30 minutes in Antigen Retrieval Solution (pH 9) (Dako), cooled down at RT and rinsed in TBS. Endogenous peroxidase activity was blocked by a 15 minute incubation in 3% H_2_O_2_. Slides were incubated in Protein Block Serum-Free (Agilent) for 30 minutes at RT, then incubated with the primary antibody diluted in Background reducing Antibody Diluent (Agilent). The primary antibody solution was washed by incubating the slides in TBS 0.04% Tween 20 and the sections were incubated with the HRP conjugated secondary antibody for 30 minutes at RT. Antigen detection was performed using the AEC Peroxidase Substrate Kit (Vector laboratories), and slides were counterstained with the Hematoxylin Solution, Harris Modified (Sigma-Aldrich). The slides were then mounted in Glycergel Mounting Media (Agilent) and imaged on a Aperio AT2 slide scanner (Leica), at a 20X magnification. To perform the subsequent staining, the coverslip was removed from the slide by incubating the slides in 60°C water, and AEC and Hematoxylin were washed away in ascending series of 50%, 70% (with 1% HCl 12N) and 100% ethanol. Sections were then re-hydrated. From that point, the staining progresses as described previously, with a shortened antigen retrieval step (10 minutes at 95°C). If two primary antibodies used consecutively were raised in the same species, an extra blocking step is performed with AffiniPure Fab Fragment Donkey anti-mouse IgG, anti-rabbit IgG or anti-rat IgG (Kackson Immuno Research), depending on the species. The following primary antibodies were used: anti-CD11b (clone EPR1344, Abcam), anti-CD4 (clone EPR19514, Abcam), anti-CD11c (clone D1V9Y, Cell Signaling), anti-CD8a (clone 4SM15, Thermo Fisher Scientific), anti-EpCAM (clone EPR20533-266, Abcam), anti-B220 (clone RA3-6B2, Thermo Fisher Scientific), anti-F4/80 (clone D2S9R, Cell Signaling), anti-AU1 tag (clone AU1, BioLegend), anti-AU5 tag (clone AU5, BioLegend), anti-Protein C tag (clone HPC4, Genscript), anti-E tag (clone 10B11, Abcam), anti-DYKDDDDK (FLAG) tag (clone L5, BioLegend), anti-HA tag (clone 6E2, Cell Signaling), anti-HSV tag (rabbit pAb, Thermo Fisher Scientific), anti-NWSHPQFEK (NWS) tag (clone 5A9F9, Genscript), anti-Ollas tag (clone L2, Thermo Fisher Scientific), anti-S tag (clone D2K2V, Cell Signaling), anti-V5 tag (clone R960-25, Thermo Fisher Scientific), anti-VSVg tag (rabbit pAb, Thermo Fisher Scientific). The following secondary antibodies were used: EnVision+ System-HRP Labelled Polymer Anti-mouse (Agilent), EnVision+ System-HRP Labelled Polymer Anti-rabbit (Agilent), ImmPRESS HRP anti-rat IgG, Mouse adsorbed (Peroxidase) Polymer Detection Kit (vector Laboratories).

#### Spatial transcriptomics

Fresh frozen samples embedded in OCT were cryosectioned and 10 µm-thick sections were placed on the Visium Tissue Optimization Slides and Visium Spatial Gene Expression Slides (10x Genomics). Tissue sections were fixed in methanol and processed using the Visium Spatial Gene Expression Kit (10x Genomics), following manufacturer’s instructions. Based on the tissue optimization time course experiment, the lung tissue sections were permeabilized for 6 minutes to maximize RNA recovery. A serial section was used for Hyperion detection of the Pro-Code epitopes (see above).

### QUANTIFICATION AND STATISTICAL ANALYSIS

#### Visualization and analysis of CyTOF data

CyTOF data was analyzed as described previously in detail (Wroblewska et al., 2018). Briefly, manual gating was performed on Cytobank and Pro-Code positive cells (based on NGFR or mCherry expression) were debarcoded using Single Cell Debarcoder (Zunder et al., 2015). To visualize debarcoding, marker intensity heatmaps were generated by taking the median of arcsine transformed intensity values, first divided by a scaling factor of 5, for each epitope tag in each debarcoded population. Transformed intensity values for each epitope tag were then rescaled from 0 to 1.

#### Image processing and visualization

MICSSS data was registered and segmented using ImageJ and QuPath, following a detailed protocol described previously (Akturk et al., 2020). Briefly, consecutive images were aligned using Image J registration tool (Linear Stack Alignment with SIFT). Alternatively, image registration was performed using a custom analysis pipeline (Chen et al, manuscript in revision). Briefly, the raw red-green-blue (RGB) 1.25x resolution image was used to generate a tissue mask. The images across all markers were then rigidly registered using the SimpleElastic k package for Python. Finally, the high resolution images (20x) were spliced into 2 mm^2^ tiles. Each set of tiles was deconvoluted to extract the hematoxylin channel, which was then registered with an affine registration (accounting for shear, scale, rotation and translational dislocation) and a bspline elastic warping (accounting for any local tissue warping or tissue damage). The vector field transformation matrix was then applied to the RGB tiles, which were concatenated to produce one final registered RGF image per marker. For each image, the signal associated with hematoxylin and the marker staining were deconvoluted in ImageJ, using the default Hematoxylin/AEC color vector. The resulting hematoxylin images from each staining were aligned to verify registration accuracy. Then, up to 10 AEC images (each corresponding to one stain) and 1 hematoxylin image were stacked, pseudo-colored and overlayed to visualize the combinatorial expression of each marker.

Cell segmentation was performed in QuPath (v. 0.2.3) on the overlayed images. Nuclear cell segmentation identified nuclei based on the hematoxylin stain, with parameters optimized for each tissue. The same segmentation parameters were used for all images of the same experiment. The cell boundaries were extrapolated by expanding the nucleus boundaries by 3 pixels (approximately 1.5 µm) in all directions. The resulting segmentation data was exported and analyzed in R and python.

#### Pro-Code Debarcoding

Pro-Codes were assigned to cells using an algorithm adapted from Zunder *et. al.* (Zunder et al., 2015). The mean nuclear pixel intensity for each epitope tag in each segmented cell was rescaled from 0 to 1 within each tissue section. Next, epitope tags were sorted on normalized signal and the three epitope tags with the highest signal were assigned as a Pro-Code for a given cell if the difference between the third and fourth highest epitope tags exceeded a given threshold which could be optimized for stringency of Pro-Code calling. In all tissue sections for this study a difference threshold of 0.2 was used. Pro-code assignments were then referenced against the vector library design to assign gRNA target genes and remove any inconsistent Pro-Code combinations.

#### Clonality

Segmented cell coordinates for Pro-code positive cells were loaded into squidpy v1.0.0 (Palla et al., 2021) for clonality and co-localization analyses. On each tissue section a neighbors graph was constructed using the 10 nearest neighbors of each cells at a maximum radius of 40 μm. Permutation based co-localization Z-scores were calculated using the nhood_enrichment function in squidpy with 1,000 permutations. This measured the relative frequency of interactions between each population of PC positive cells and all other PC populations by taking the Z-score of the observed number of interactions between two PC populations within a permuted background generated by swapping PC labels. Group degree centrality (fraction of non-homotypic neighbors) and average local clustering coefficient graph metrics were calculated for cells positive for each Pro-Code on each tissue section using the centrality_scores function in squidpy on the neighbors graph mentioned previously. Graph metrics were only calculated for Pro-codes with greater than 20 assigned cells on a tissue section. Kernel densities for abundant Pro-Code populations in 4T1 primary tumors were estimated using the kde2d function from the MASS (Venables and Ripley, 2002) package in R that used a bivariate normal kernel with normal distribution approximation to determine bandwidth.

#### Clustering and lesion definition

KP tumor cells tend to form clonal lesions as we observed based on Pro-code tagging. To sub-cluster Pro-code marked KP cells into clonal lesions and remove stray outlier cells we used the density based spatial clustering of applications with noise (DBSCAN) (Hahsler et al., 2019) algorithm separately for each Pro-code within each tissue section that was assigned to at least 10 cells. For DBSCAN, an epsilon neighborhood size of 200 μm and minimum number of cells in the epsilon region set to 10 were used. Boundaries were drawn around clonal lesions using alphahull, a derivative of the convex hull algorithm that allows for concave boundaries, using the ashape function of the alphahull package (Pateiro-López and Rodríoguez-Casal, 2010). The alpha value, a tuning parameter where the higher the value the closer approximation to the convex hull, was initialized at 80 for each clonal lesion and borders were drawn iteratively increasing this value by 10 if a closed loop was not formed. Defined lesions were then exported to QuPath and subjected to manual curation based on image inspection to ensure quality and remove spurious associations. Lesion size was calculated by the pixel coordinate area of the resulting boundary polygons. Only tumors greater than 25,000 μm^2^ were kept for subsequent analysis.

#### Tumor immune cell composition

Segmented cells were called positive for each phenotypic marker by using standardized intensity cutoffs manually inferred from signal to background inspection of images as well as intensity histograms. Mean pixel intensities for CD4, CD8, B220, CD11b, CD11c, and F480 were determined using mean intensity across the whole segmented cell. EPCAM values for each tumor lesion were quantified as the mean cellular pixel intensity of all segmented cells within the tumor area. The composition of cells positive for each phenotypic marker was quantified in defined tumor boundaries using windows of the full tumor area (100% of border), along with the tumor core (inner 70%) and periphery (±10% from border). The frequency of phenotypic marker positive cells was normalized per unit area in each tumor window (cells per μm^2^). The significance of differences in normalized immune composition scores for each marker in the myeloid and lymphoid marker panels was assessed by a Wilcoxon rank-sum test between F8 (off-target control) tumor lesions and tumors for each gene perturbation that had greater than 20 identified tumor lesions. P values for each panel were adjusted for multiple testing by Benjamini-Hochberg correction (Benjamini and Hochberg, 1995). Composition scores were visualized in radial plots, where the height of the corresponding bar for each marker was the difference in median normalized composition scores between F8 control tumors and tumors of a given gene perturbation, with the shading of bars corresponding to the signed −log_10_ adjusted p value of the Wilcoxon rank-sum test described above. A distance threshold of 75 μm between curated tumor lesion boundaries was used to identify neighboring lesions for the analysis in figure 4F.

#### Histology associations

Significance of relationships between gene perturbation and pathology scores described above were defined with a Chi-squared test using the chisq.test function in R for each annotation category separately (location, degree of differentiation, stroma, pattern). Input for these tests were contingency tables of gRNA target and categorical annotations specific to each category, with 1,753 annotated tumors with defined Pro-codes as input. Associations between specific annotations and gene perturbations were assessed by the standardized residuals of the Chi-squared test. Tumor sizes were determined from curated tumor boundaries after debarcoding in the lymphoid marker-stained sections.

#### Spatial Transcriptomics analysis

The 10x Genomics Spaceranger mkfastq (v. 1.2.1) pipeline was utilized to demultiplex raw base call files into FASTQ format. The FASTQ files along with a microscopy brightfield image stained with hematoxylin and eosin (H&E), was used by Spaceranger count (v 1.2.1) to perform alignment to a modified mm10 genome, tissue detection, fiducial detection, and barcode/UMI counting. The modified mm10 genome contained additional sequences for the WPRE sequence of the Pro-Code-bearing transcript, to identify Pro-Code expressing spots. The pipeline used the Visium spatial barcodes to generate feature-spot matrices. Spots deemed off-tissue by visual inspection were annotated in the Loupe browser and removed from subsequent analysis.

Analysis of spatial transcriptomic data was conducted primarily using Seurat v 4.0.3 (Hao et al., 2021). To define tumor versus normal spots in the entirety of each tissue section first library size normalization was performed across all spots within each section, dividing the feature counts by the total number of reads for each spot and then multiplying by a scaling factor of 10,000 followed by natural log transformation using the log1p function in R. Highly variable genes (HVG) for each tissue section were selected using the FindVariableFeatures function in Seurat with the ‘vst’ setting for 8,000 features. Feature information was then scaled and centered followed by principle component analysis (PCA) for dimensionality reduction. Coordinates of the top 40 principle components were input into the kmeans function in R with k = 2 to cluster tumor versus normal spots on each slide. To create a conserved tumor vs normal expression signature, a Wilcoxon rank-sum differential expression test between tumor and normal clusters on each slide was performed, genes that were differentially expressed (Bonferroni adjusted p < 0.1) on at least three out of four slides and with a consistent direction of effect were included in the conserved signature (908 genes). Raw WPRE unique molecular identifier (UMI) counts (corresponding to Pro-Code transcripts) within spots for each cluster were compared to confirm the clustering distinction and the WPRE read count distribution within tumor spots across sections confirmed similar expression. Subsequently, tumor spots were defined as spots with raw WPRE UMI counts greater than or equal to 4, the mode of the first quantile across kmeans defined tumor spots in each section.

For tumor specific clustering, WPRE defined tumor spots within each section were renormalized and processed in the same way as for the tumor versus normal comparison except 5,000 HVGs were used for input into PCA and the top 20 principle components were used for Leiden clustering (Traag et al., 2019) of tumor specific spots. Tumor spots on each Visium section were clustered separately. Leiden spot clusters were annotated based on spatial location and overlap with regions specific for certain Pro-codes from parallel Hyperion imaging. Tumors were grouped into clusters segregating specifically for certain gene knockouts (Tgfbr2, Ifngr2, and Jak2) as well as non-specific tumor types (named KP followed by and underscore and the slide index, then a dash for cluster number) and spots appearing to correspond to locations on the periphery of the tumor. Following slide specific clustering, all tumor spots with their corresponding annotations were merged together and jointly normalized using SCTransform (Hafemeister and Satija, 2019) to compare across tissue sections. UMAP (Becht et al., 2019) dimensionality reduction using all tumor spots was done using the top 20 principle components derived from 3,000 HVGs. Hierarchical clustering of tumor cluster mean spot expression was performed using a Pearson correlation distance and average agglomeration of mean spot scaled Pearson residuals for each feature calculated in the SCTransform algorithm. Tumor cluster specific gene signatures were calculated by a Wilcoxon rank-sum test using normalized expression values between canonical KP tumor lesions (clusters KP_2-1, KP_4-1, KP_1-2, KP_4-4, KP_4-2, KP_4-3, KP_2-3, KP_3-2, KP_3-3) and the periphery, Ifngr2, Jak2, and Tgfbr2 knockout clusters. Only genes with at least a 0.25 log_2_ fold-change between the groups and UMI count detection in at least 10% of spots for either cluster were input into the differential expression test. Significant differentially expressed genes (DEG) had a Bonferroni adjusted P value < 0.01. Fgsea (Korotkevich et al., 2016) with the average log_2_ fold-change of DEGs was used to determine gene set enrichments within gene ontology (GO) terms and Tgfb response sets (TBRS) in fibroblasts, endothelial cells, macrophages, and T cells defined in Calon *et al* (Calon et al., 2012). Human gene symbols from TBRS gene sets were converted to mouse by Ensembl BioMart (http://www.ensembl.org/biomart/martview/) and only one-to-one orthologues were used. Gene modules relating to Ifngr2 and Tgfbr2 targeted tumors were generated by hierarchical clustering (Pearson correlation distance and average agglomeration, tree cut at 7 clusters) of all DEGs for either population with an average log_2_ fold-change greater than 0.5. As input, mean log normalized expression values for each tumor cluster except Jak2_1 and KP_3-1 were used.

#### Statistical analysis

Statistical analysis for this study was performed using the R and python programming languages using the packages and methods described above. Gene perturbations with a low number of detected lesions were excluded from differential immune infiltrate calculations.

#### Visualization

Figures for the analysis of Imaging or genomic data were created in QuPath, ImageJ, or R. The geom_density_2d function from ggplot2 (Wickham, 2009) was used to visualize kernel density estimates of Pro-Code tagged cells. Plots of Visium spots overlaid onto tissue images were generated using Seurat.

#### Data availability

The datasets generated in the current study are available from the corresponding author on reasonable request. The memPC library is scheduled for distribution by Addgene by August 2021. nPC will also be available from Addgene but for immediate request Authors can be contacted.

#### Code availability

Custom code used in the analysis of Perturb-map imaging data can be found at https://github.com/srose89/PERTURB-map.

---

**Supplementary Figure 1, related to Figure 1 and 2.**
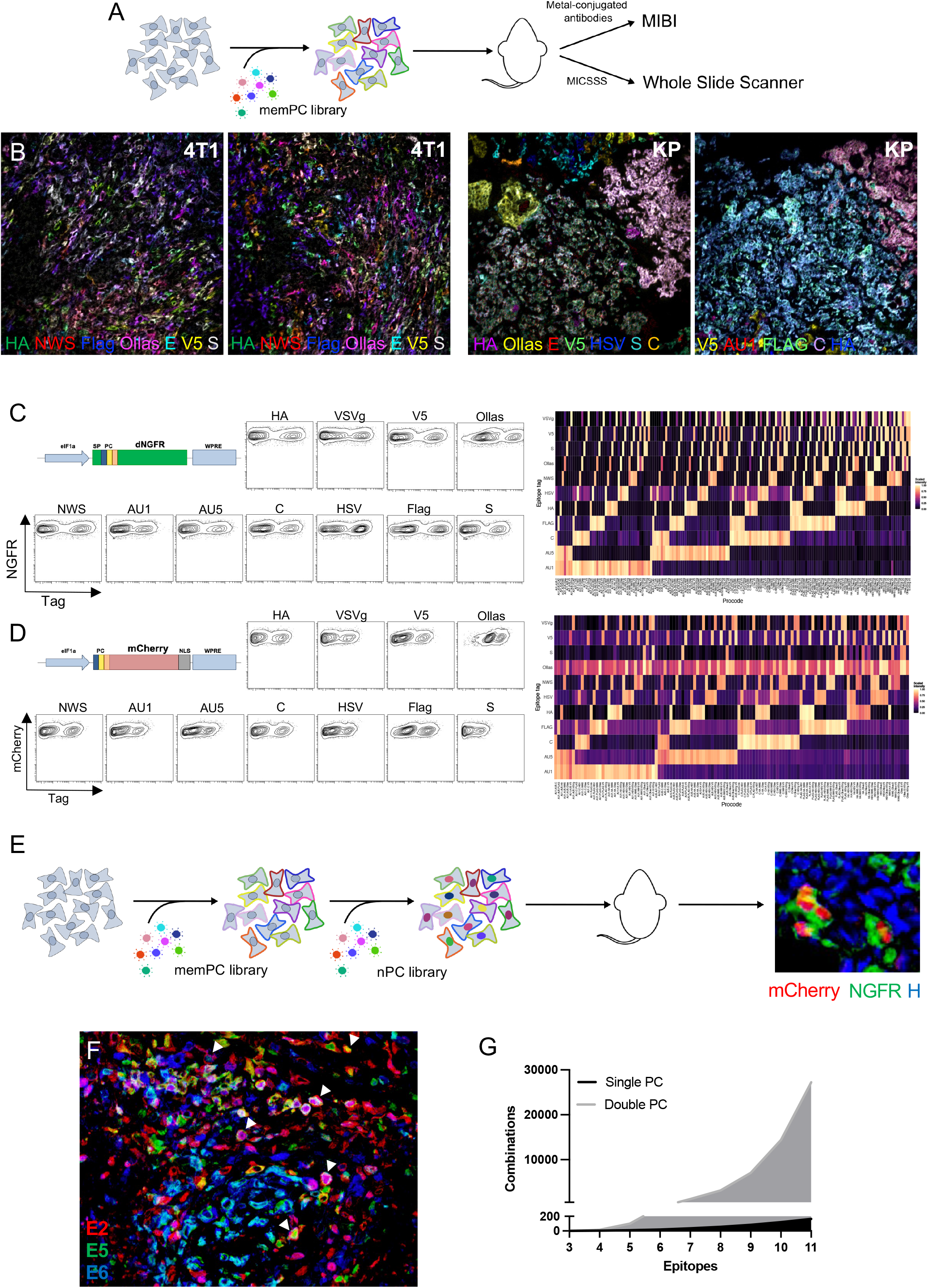
dNGFR Pro-Codes (memPC) and nuclear Pro-Codes (nPC) are detected by multiplexed imaging and mass cytometry. (**A**) Schematic of the experimental setup for multiplex imaging of the Pro-Codes in breast and lung tumors. 4T1 or KP cells were transduced with a library of 120 memPC vectors and injected into BALB/c or C57BL/6 mice. Tumor-bearing organs were harvested and either fresh frozen (for MIBI) or formalin fixed paraffin embedded (for MICSSS). For MICSSS, tissue sections were stained consecutively with anti-epitope tag antibody, and imaged on a whole slide scanner. For MIBI analysis, tissue sections were stained with a cocktail of anti-epitope tag metal-conjugated antibodies and imaged on the MIBI instrument. (**B**) Multiplex Ion Beam Imager (MIBI) analysis of Pro-Codes in 4T1 breast and KP lung tumors. Shown are overlaid images from MIBI acquisition. Selected tags are shown, color-coded as indicated. Representative of 20 field of views acquired across 3 independent experiments (4T1) or 6 fields of view from one experiment (KP). (**C**) CyTOF analysis of membrane-bound Pro-Code (memPC) library distribution. 293T cells were transduced with a library of 165 memPC and analyzed by CyTOF. (Left) Individual epitope tag staining. (Right) Heatmap representing the scaled (from 0 to 1) median intensity of each epitope tag for each memPC following debarcoding. (**D**) CyTOF analysis of nuclear localizing Pro-Code (nPC) library distribution. 293T cells were transduced with a library of 165 nPC and analyzed by CyTOF. (Left) Individual epitope tag staining. (Right) Heatmap representing the scaled (from 0 to 1) median intensity of each epitope tag for each nPC following debarcoding. (**E**) Schematic of the experimental setup for compound Pro-Code library usage. 4T1 cells were transduced with a library of 56 memPC, sorted based on dNGFR expression, transduced with a second library of 56 nPC and sorted based on mCherry expression. Cells were then injected into BALB/c mice and the resulting tumors were stained by MICSSS for each of the Pro-Code epitopes. Representative image of Pro-Code positive cells in the tumor, showing nuclear localization of mCherry (nPC) and membrane localization of dNGFR (memPC). (**F**) Multiplex imaging analysis of compound Pro-Code library expression in a 4T1 breast tumor. Shown is an overlaid image of a 4T1 tumor expressing both memPC and nPC libraries (established as described above). The staining corresponding to 3 epitope tags was overlaid and pseudocolored. Arrowheads point to example of cells expressing a different epitope tag combination at the membrane (memPC) and nucleus (nPC). Representative image from 5 mice is shown. (**G**) Projection of the number of possible combinations that can be tracked in vivo using a single Pro-Code LV library or double-transduced cells (memPC and nPC). Compounding two libraries with 8 epitope tags can achieve 3,136 combinations. Compounding two libraries with 11 epitopes tags (the number we have validated for in situ detection) can achieve 27,225 combinations.

**Supplementary Figure 2, related to Figure 2.**
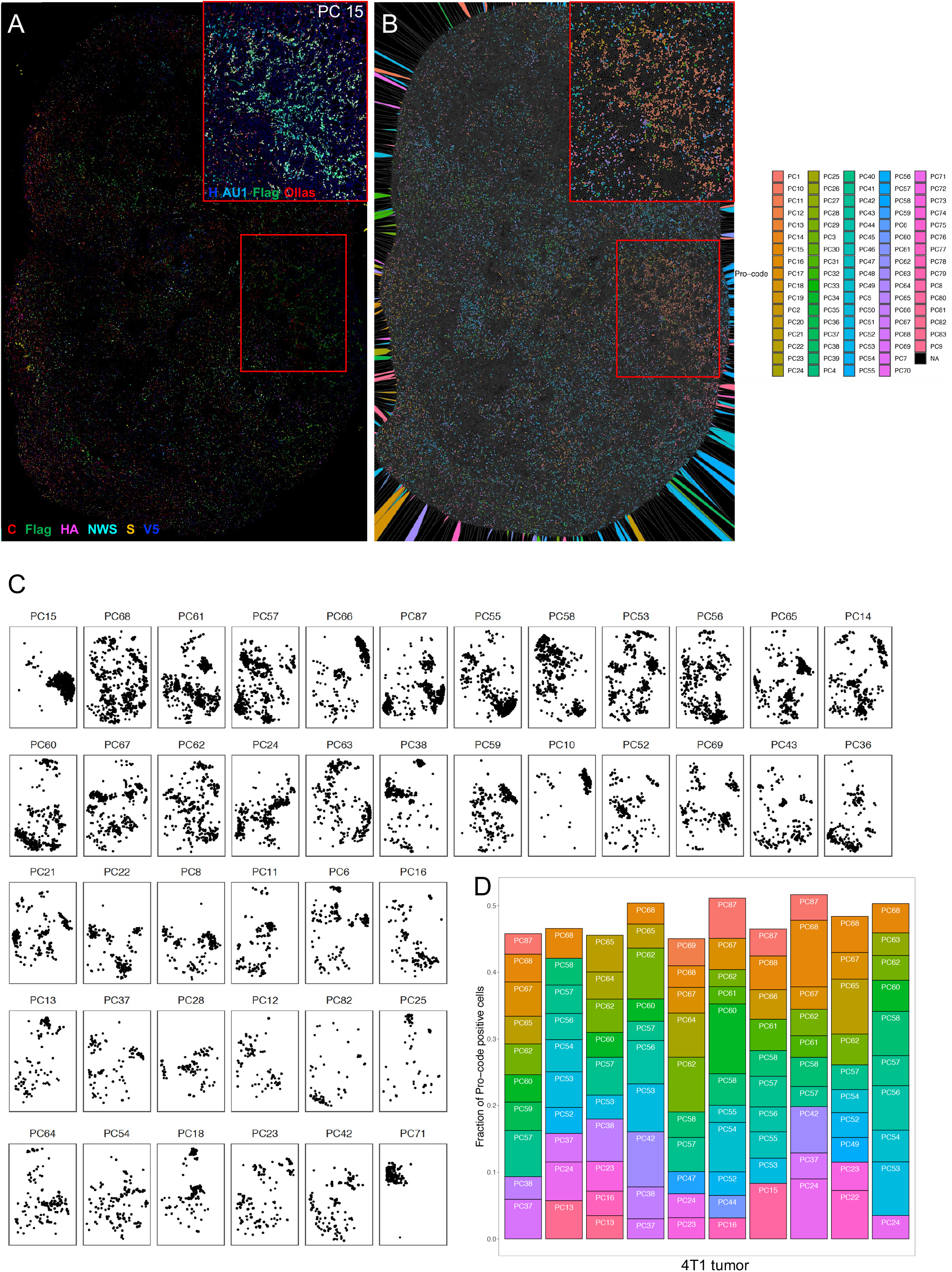
Cell tracking of 4T1 tumors in vivo. (**A**) 4T1 cells were transduced with a nPC library (120 combinations, 10 epitopes) and injected in the mammary fat pad of female BALB/c mice. The tumors were harvested 2 weeks later and consecutively stained with antibodies specific for the 10 epitope tags by MICSSS. Shown is a representative image overlaying 6 epitope tags (as indicated), as well as a zoomed in region of interest (ROI) highlighting PC 15 (composed of AU1, Flag and Ollas). Image representative of 16 tumors, across 2 different experiments with different nPC libraries (same cohorts as in Figure 2). (**B**) Voronoi plot of the same image as in (A), after Pro-Code debarcoding using the difference in scaled intensity between the third and fourth most intense epitope tags for each segmented cell. Enlarged ROI highlights PC 15 (in orange). (**C**) Individual plots representing the coordinates of Pro-Code+ cells for the 42 most abundant Pro-Code population from (A). Each dot corresponds to a Pro-Code+ cell, and the 2D map is reconstituted based on each cell’s XY coordinates. (**D**) Analysis of the PC composition of each tumor. For each tumor, the relative abundance if the 10 most frequent nPC detected in a tissue section was plotted. Shown are 10 tumors from one experiment, representative of 2 experiments.

**Supplementary Figure 3, related to Figure 3.**
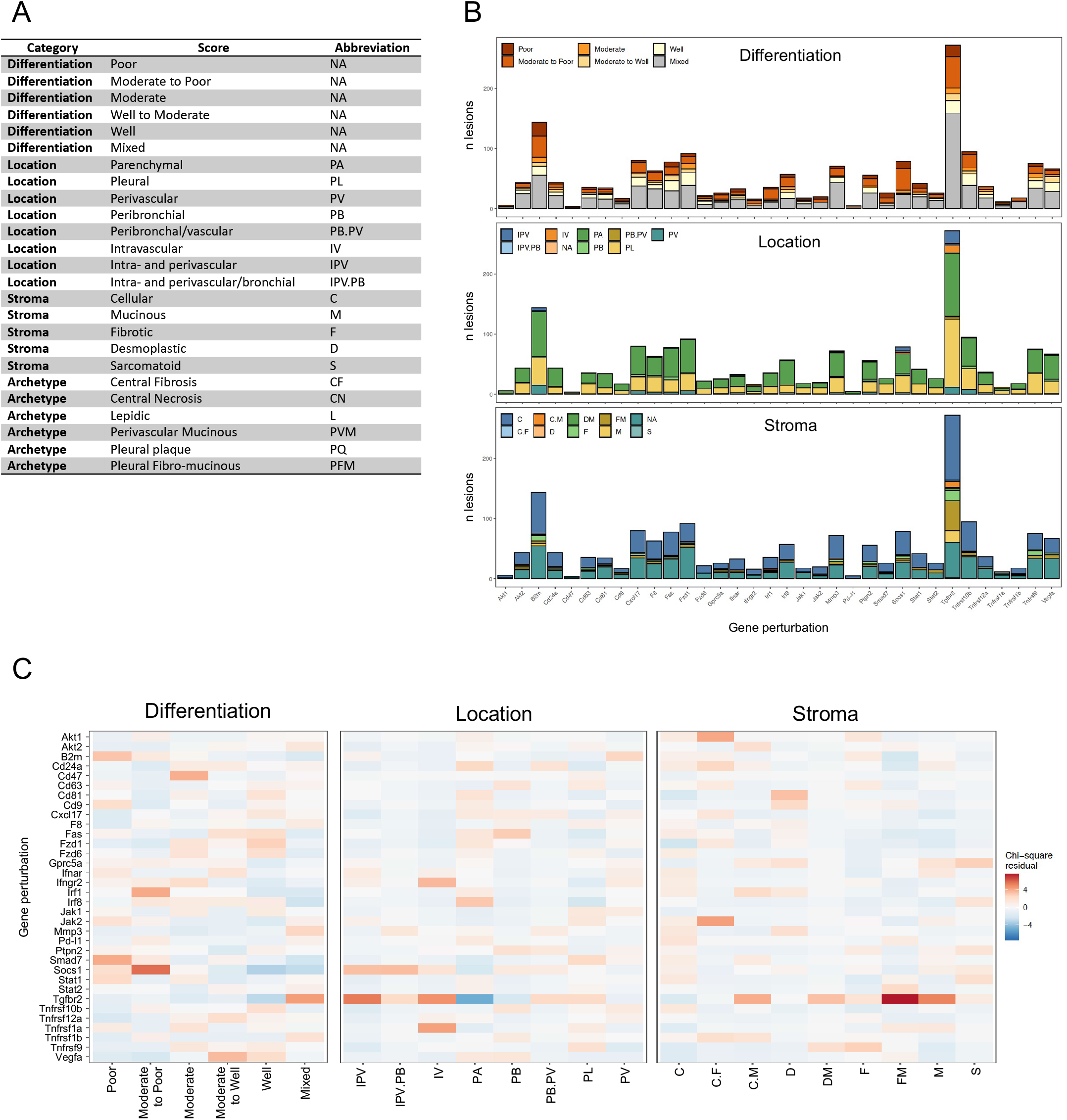
Perturb-map identifies the contribution of gene perturbation to the architecture of KP tumor lesions. (**A**) Table of the categories used by a trained pathologist to score KP tumor lesions. For each tumor, the differentiation degree was evaluated (from well to poor), as well as the lesion location within the tissue and the composition of the tumor associated stroma. Independently, tumor archetype were identified across samples based on a combination of criteria (as described in the Results). (**B**) Overview of the repartition of tumors within each scoring category related to differentiation, location and stroma composition. Each tumor lesion was scored independently on a serial section stained with H&E by a pathologist blinded to the perturbations. Each column corresponds to the gene perturbation. The bars are colored as indicated. (**C**) Quantification of associations between gene perturbations and histopathological features. The heatmaps show the standardized residuals of a Chi-squared test for each gene perturbation and each scoring category. (Chi-squared p value, Location = 0.02849, Stroma = 5.05×10^−5^, Differentiation = 8.933×10^−9^)

**Supplementary Figure 4, related to Figure 5.**
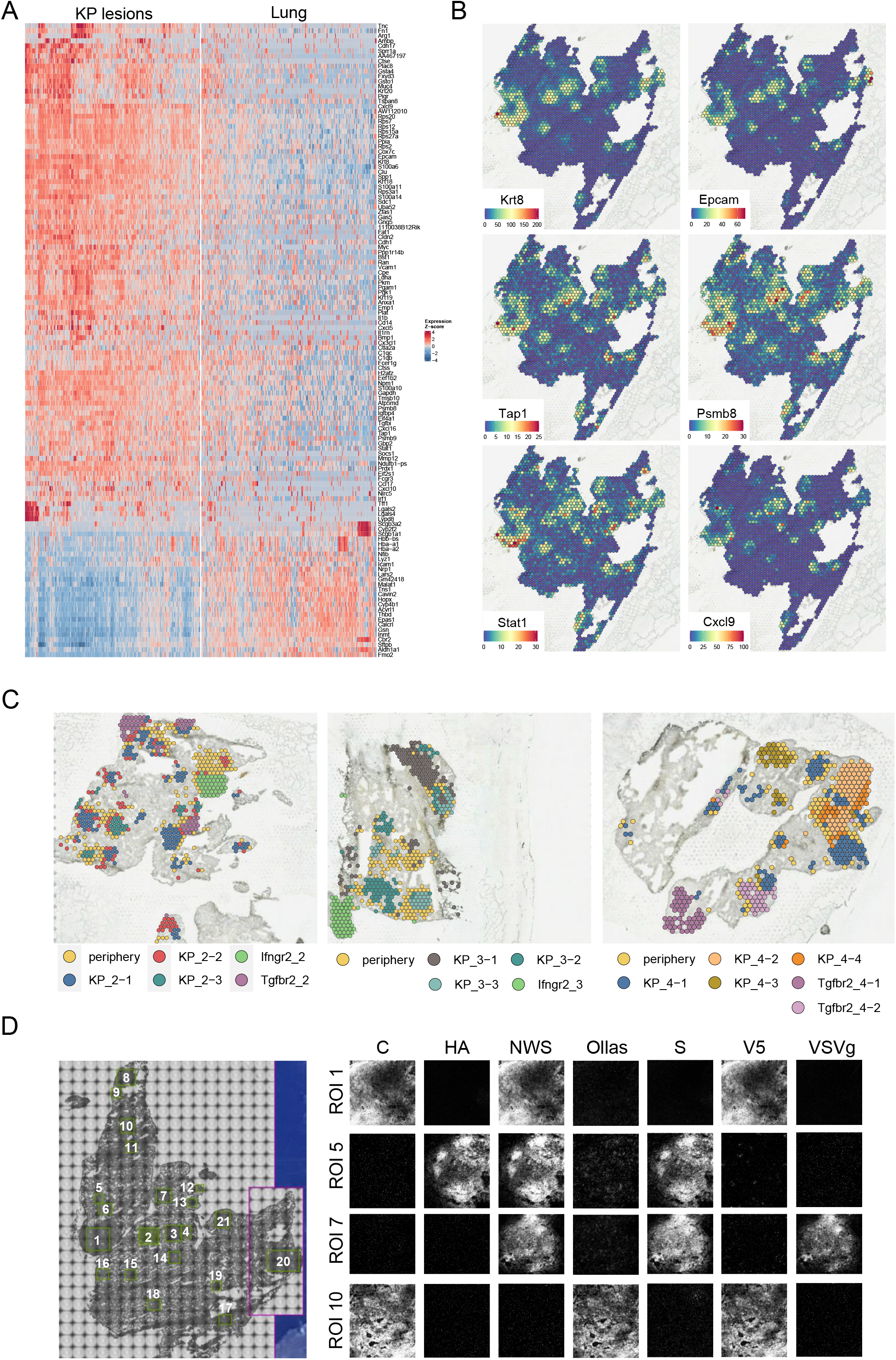
Perturb-map can be coupled with Spatial Transcriptomics. (**A**) Gene signature of KP lesions compared to normal lungs. A representative sample of 250 spots for each cluster is shown. Color is shaded by row Z-score of log normalized counts, with genes and spots arranged by hierarchical clustering using Pearson correlation distance and average agglomeration. Displayed are the top 75 and 25 genes with the highest log_2_ fold-change in the tumor or non-involved lung clusters, respectively, in addition to IFNγ regulated genes within the signature. (**B**) UMI counts of selected tumor-specific or IFNγ pathway transcripts. (**C**) Leiden defined clusters of Pro-Code^+^ spots (>= 4 WPRE UMIs) in Visium profiled tissue sections. 2,006 total tumor spots were analyzed across 4 tissue sections, with spots clustered separately for each section. (**D**) For each sample, we generated two serial sections, to be used respectively for Visium (section 1) and for identification of the gene perturbation within tumor lesions (section 2). Section 2 was stained with metal-conjugated antibodies specific to each epitope tag and imaged by imaging mass spectrometry (Hyperion). (Left) Tissue image, displaying the region of interests (ROIs) selected for ablation (green rectangles). (Right) Signal associated to each epitope tag in selected ROIs. Each ROI was positive for a triplet combination of epitope tag, identifying the Pro-Code and associated gene perturbation. In total, we imaged 77 ROIs across 4 tissue sections (corresponding to 4 lobes of lung).

